# Summarizing Evolutionary Trajectories from Phylogenetic Character Maps of Discrete Traits

**DOI:** 10.64898/2026.06.14.732171

**Authors:** Sean W. McHugh, Michael J. Landis

## Abstract

When reconstructing phylogenetic character histories, biologists aim to identify distinct evolutionary trajectories, or paths of character state evolution. However, biologists typically wish to summarize the information representing large numbers of potential character histories for a single phylogeny. For discrete characters, few approaches exist for summarizing the number of unique evolutionary trajectories beyond the frequency of specific events (i.e., state transition types) or the time lineages spend in each state. Here, we introduce a framework for summarizing the evolutionary trajectories of discrete character histories by compressing them into trajectory trees, where branches represent unique character-evolution pathways rather than lineages. This framework includes a novel compressed tree representation, called a scenario tree, that retains temporal information, ensuring that each root-to-tip path represents a unique, temporally explicit evolutionary trajectory. We describe and apply several approaches to summarize phylogenetic trees into transition trees. We include visual summaries – such as consensus trajectory trees, trajectory tree tanglegrams, and“trajectory-through-time plots” – to compare how unique evolutionary trajectories accumulate across lineages and state transitions. We also include quantitative summaries, such as the time spent in unique evolutionary trajectories and the number of transitions that follow unique character-state transitions. We use our new trajectory-wise summaries to evaluate the adequacy of commonly used continuous-time Markov models of character evolution, which are memoryless and consider only the rates between pairs of states. We conducted multiple simulation-based experiments demonstrating the utility of our novel trajectory-wise approaches. We also apply our new trajectory-wise approaches to Greater Antillean Anolis lizard biogeography and ecomorph evolution, and find that Anoles evolved along considerably more unique evolutionary trajectories than expected under simulations of our best-fitting character evolution model. The number of unique evolutionary paths accumulated in an “early burst” pattern relative to simulated trajectories, with this burst being more intense than expected across all character state transition events.

## Introduction

Biologists commonly seek historical explanations for the origin and disparification of key characters of evolutionary significance. Common examples include a lineage that attained a key innovation (Donoghue 2005; Hunter 1998; Erwin 2015; Miller and Stroud 2021), a pathogen lineage acquiring novel amino acid substitutions to become more virulent (May and Rannala 2024), or a somatic cell lineage gaining driver mutations to become cancerous (Vendramin et al. 2021; Mitchell et al. 2022). To explain these historical scenarios, biologists often test hypothesized narratives of character evolution by comparing phylogenetic models and interpreting patterns of reconstructed ancestral states (Greene 2017).

When reconstructing ancestral character state histories, biologists often wish to identify distinct *evolutionary trajectories*, which we define as paths of character state evolution with unique sequences and/or timings of state changes. For instance, a biologist studying a clade of lizards may be interested in summarizing how different lineages became specialized to a particular microhabitat, such as arboreality, and gained specialized adaptive traits, such as toepads (Miller and Stroud 2021; Gamble et al. 2012) or claws (Yuan et al. 2019; Crandell et al. 2014; Baeckens et al. 2020) for climbing. The exact sequences and timings of transitions into and out of these states may be the same or vary across the phylogeny. Shared ancestry and trait conservatism may result in phylogenetic patterns where an ancestral lineage occupies a certain microhabitat, gains an adaptation to that habitat, and then diversifies into many species, with no further changes to either trait; these species would all share one evolutionary trajectory. A cartoon of an analogous character evolution pattern for three states (“green”, “orange”, and “blue”) is shown in Figure 1a-b. While in the first character history (row 1) every species has a distinct root-tip character history, the other two histories (rows 2 and 3) show instances where multiple species (row 2: spp. 1 and 3; row 3: spp. 2 and 3) share a conserved evolutionary trajectory from a common ancestor.

**Figure 1:**
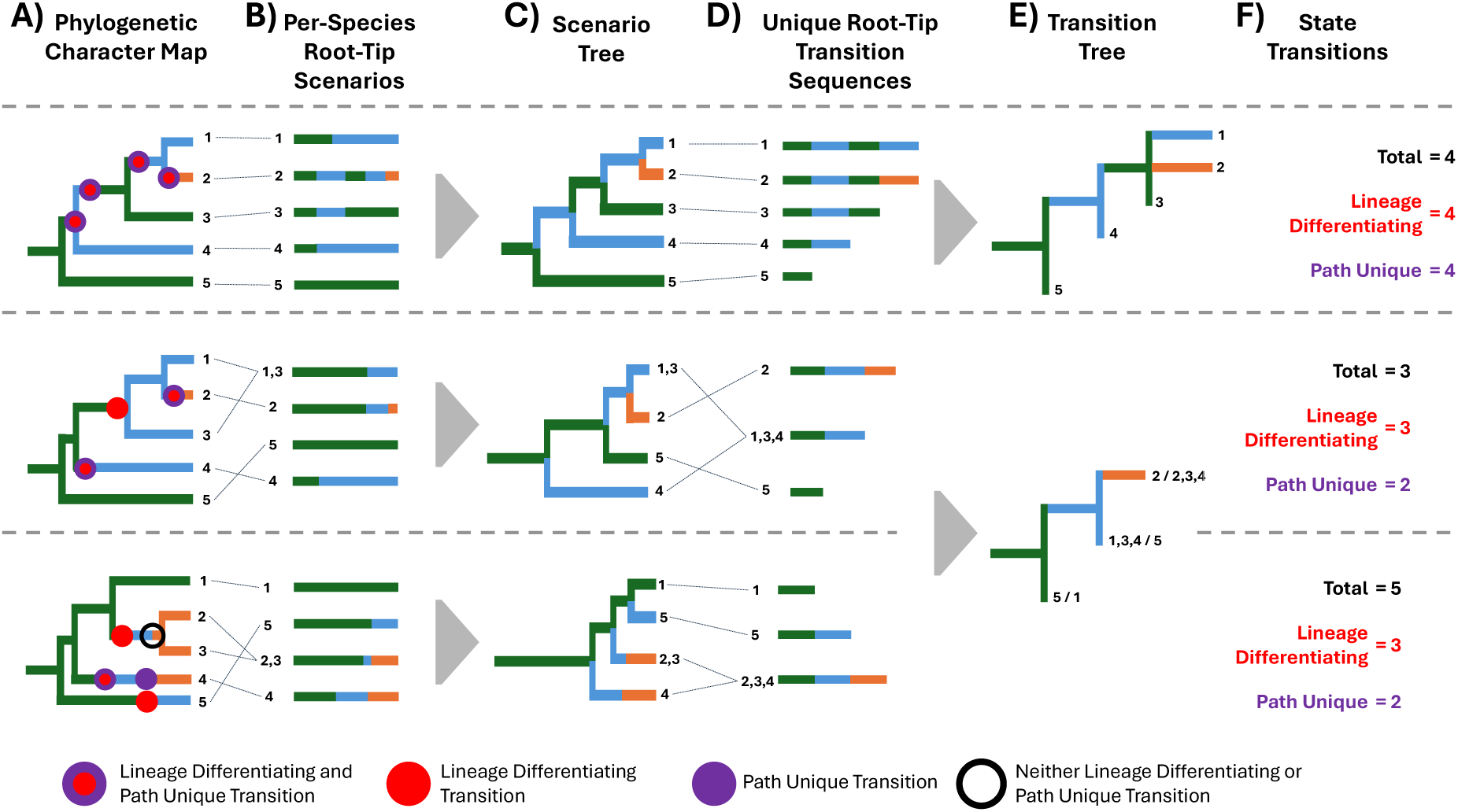
Cartoon of the relationship between phylogenetic character histories, evolutionary tra-jectories, and summary statistics for a system with five species (1-5), three character states (green, blue, and orange), across three separate evolutionary histories (one per row). Each history has six panels. A) The phylogenetic character history. B) The unique root-tip character history scenarios across species (shown in units of time), with dotted lines showing which species correspond to which scenarios. C) The phylogenetic character history compressed to a scenario tree, with each tip representing a unique scenario, with branching occurring when two lineages diverge in character history. D) The unique character history transition sequences (in units of number of character state transitions). E) The phylogenetic history compressed into a transition tree, with each tip representing a unique state transition sequence. F) The number of state transitions of each type, including: total number of state transitions, the number of lineage-differentiating transitions (i.e., transitions that result in scenario tree splitting), and path unique transitions (i.e., transitions resulting in transition tree branching). The first history shows an instance in which each species follows a separate scenario and transition sequence. The other two histories (second and third rows) show instances in which more than one species evolves under the same scenario (A and C in the second row, and B and C in the third row). Importantly, while the timing of state transitions differs across histories two and three, the two histories follow the same transition tree.

The timing and/or the sequence of events can cause trajectories to vary among lineages. For example, some may only transition into a given microhabitat only after gaining geographical access to it, while others may need to acquire an adaptation, such as toepads, beforehand. Although both species evolutionarily arrived to the same microhabitat with the same adaptations, their trajectories could differ in terms of transition sequence (Figure 1b). Yet other species could also follow the same transition sequence, leading to the same microhabitat, but with different timings. One species may shift early in its trajectory, while the other species shifts much later. These trajectories would differ only due to differences in the timing of transitions (Figure 1b-e, rows 2 and 3). Across a large phylogeny, such as one including all lizards, many diverse trajectories with variable transition sequences and timings may result in species occupying similar habitats.

Identifying patterns among trajectories is critical to answering important biological questions, including: Which evolutionary events or event sequences tend to result in the greatest evolutionary “success” in terms of either species diversification or trait disparification (Hunter 1998; Donoghue 2005; Hautmann 2020)? Do trajectories diverge before or after major historical events such as climatic or geographic shifts (Berv et al. 2024)? How prevalent are convergent or parallel trajectories (Esquerŕe and Scott Keogh 2016; Monnet et al. 2011; Bonett et al. 2022)?

While substantial progress has been made in developing biologically realistic phylogenetic models for reconstructing discrete character histories (Tarasov et al. 2019; Beaulieu and O’Meara 2016; Boyko and Beaulieu 2023), generating the evolutionary trajectories that represent these histories remains challenging. Existing summaries of discrete character histories—particularly those derived from stochastic character mapping (Huelsenbeck et al. 2003)—tend to reduce complex evolutionary histories to event-wise or state-wise statistics, such as state transition and state-time frequencies (Farris 1989; Archie and Felsenstein 1993; Lemey et al. 2012; Rodrigue et al. 2008; Laurin-Lemay and Rodrigue 2025). While informative in many ways, and often still underutilized (Lemey et al. 2012; Laurin-Lemay and Rodrigue 2025), they are summaries that do not fully capture the diversity of evolutionary trajectories. Identical numbers of transitions can arise from histories composed of many redundant trajectories (Griffing et al. 2022) or from histories that traverse diverse and distinct sequences of character change (Williams et al. 2013; Hagman and Ord 2016; Twomey et al. 2023). Depending on the structure of the transition rates throughout the phylogeny, the degree of lineage differentiation can vary considerably. When state transitions occur in one of many lineages following a conserved history, those lineages become differentiated with one possessing a now “derived” state relative to the “ancestral state” (Hoyal Cuthill 2015). Such lineage differentiation does not occur when transitions accumulate on single long branches. Additionally, the shared history (or “tree”) of evolutionary trajectories and its corresponding phylogeny can closely match or differ in topology, depending on how closely lineage divergence and subsequent state divergence in daughter lineages match. Identifying these important patterns requires an explicit, trajectory-level view of character evolution that preserves the sequence, timing, and phylogenetic context of discrete state transitions.

Identifying important trajectory-wise patterns becomes challenging as the phylogeny size and the number of tip states increase. Biologists frequently summarize the evolutionary histories of many important discrete characters over large clades of hundreds to thousands of taxa (Sauquet et al. 2017; Blaimer et al. 2023). Such important characters are often correlated with suites of other traits, including life history, ecological mode, habitat affinity, diet, and large sets of underlying genetic changes. Few quantitative approaches can summarize such large character histories beyond the state-wise and transition-wise statistics discussed earlier. One of the few approaches developed to summarize such high-dimensional discrete state spaces encodes character-state paths as trajectories on a hypercube (Williams et al. 2013; Greenbury et al. 2020; Aga et al. 2024). While hypercube approaches can effectively summarize many details about evolutionary trajectories, they require a known starting root state and a maximally derived terminal state (e.g. a set of binary alleles with the start being all “wild type” and the terminals being all “mutant”), they only consider the state transition sequence, but not the timings of the transitions within the sequence, and they present trajectories as networks where separate trajectories that arrive on the same state in the same sequence step merge to the same node.

In practice, biologists often identify evolutionary trajectories by constructing a tree-like representation of possible character state transitions, which we refer to as a trajectory tree. The history of such character-specific trees is extensive. It predates many phylogenetic comparative methods (Zimmermann 1934), originating in the organization of taxa into major groups based on the relationships between homologous structures (Mickevich and Weller 1990; Zimmermann 1952; Donoghue and Kadereit 1992), such that the character trajectories informed the phylogenetic relationships among taxa. Current approaches follow a reverse pattern of inferring character trajectories from ancestral state reconstructions mapped onto previously inferred phylogenies (Pagel 1999). Condensing character evolution into tree-like patterns typically involves “compressing” (Ishikawa et al. 2019) ancestral state reconstructions on phylogenies, in which tips represent unique character-state sequences and branches represent conserved segments of those sequences (Chevenet et al. 2013, 2019; Ishikawa et al. 2019; Chevenet et al. 2024). Transition trees provide a particularly concise representation of the diversity and shared ancestry underlying the evolutionary trajectories implied by a character map. This compression, however, entails a loss of temporal information. Transition tree branches are typically scaled by the number of transitions rather than evolutionary time, obscuring when transitions occurred and how often similar trajectories arose independently (Chevenet et al. 2013). As a result, lineages that share the same sequence of states but differ in timing may be collapsed into a single trajectory, limiting the ability to distinguish how many steps of a transition sequence are conserved or convergent amongst lineages. For example, in Figure 1, the bottom character history shows many species following a “green”-“blue”-“orange” transition sequence. From the transition tree, it is clear that three species followed this transition sequence (Figure 1E), but two species (2,3) share the same trajectory from the same common ancestor, while a separate species (4) independently transitioned from “green”-“blue”-“orange”. Moreover, transition trees are most often reported as single summary objects derived from maximum-likelihood ancestral state reconstructions at nodes. Still, they are rarely summarized as numerical quantities, as is necessary for extracting information from distributions of stochastic character maps.

There is a need for additional trajectory-level representations that preserve both the sequence and timing of character change, and for quantitative summaries of categorical trajectories that can be quantified across distributions of inferred histories. Quantitative summaries are especially useful as they can be integrated into many of our existing approaches for evaluating model adequacy and hypothesis testing (Schwery et al. 2023; Laurin-Lemay and Rodrigue 2025). Most phylogenetic models of character evolution are memoryless and only consider rates of character evolution given the current state a species possesses, rather than the order by which a species previously acquired states leading to its current state (Pagel 1999; Lewis 2001; Goldberg and Foo 2020). Given this memoryless property, an important question when applying character evolution models is whether the patterns in evolutionary trajectories inferred from our ancestral state reconstructions fall within the expected ranges predicted by our best-fit models.

Here, we present a framework for trajectory-wise summaries of discrete character histories that builds upon previous approaches. Our framework includes descriptions of the types of trajectories we can identify from discrete character maps and the subsequent tree structures we use to represent the trajectories that describe a clade’s evolutionary history. We introduce a new compressed tree representation, called a scenario tree, which retains temporal information about character histories. When compressing a phylogeny into a scenario tree, lineages are merged only when they share the same transition sequence and remain temporally synchronized, so that each root-to-tip path represents a unique, temporally explicit evolutionary trajectory. Using trajectory trees, including both scenario and transition trees, we develop additional visual summaries inspired by widely accepted approaches for summarizing patterns of lineage evolution. We also accompany our new visual summaries of trajectory trees with new statistical tests. We use quantitative information extracted from trajectory trees to evaluate whether simple phylogenetic models of character evolution can adequately describe patterns of unique trajectory-wise character evolution. We conducted multiple simulation-based experiments demonstrating the utility of our novel trajectory-wise approaches.

Finally, we applied our trajectory-wise approaches to summarize the evolutionary history of Caribbean *Anolis* lizard biogeography and ecomorph evolution. *Anolis* has successfully colonized many islands in the Greater Antilles, and radiated to occupy many microhabitats among these regions, far more so than one would expect for a clade of this size (Losos 2011; Stroud and Losos 2020; Muñoz et al. 2023). While biologists have studied the extent of geographic and niche space that *Anolis* lizards occupy, most studies of *Anolis* evolutionary trajectories have focused on quantitative morphological characters (Losos 1992; Huie et al. 2021; Losos et al. 2006; Losos 1990). Here, using stochastic character maps and our new methods to represent evolutionary trajectories, we describe the amount of unique categorical evolution in ecomorph evolution, historical biogeography, and their joint history. Specifically, we aim to (1) identify the dominant scenarios and transition sequences that Greater Antillean *Anolis* lineage may have followed; (2) characterize the degree of congruence between evolutionary trajectories and lineage diversification; (3) test whether the accumulation of evolutionary trajectories falls within expected ranges predicted by a simple inference model.

## Methods

### Formalizing Evolutionary Trajectories

We use the term *evolutionary trajectory* to refer to a root–to–tip sequence of discrete character states traced along a phylogeny (Table 1). A trajectory describes a series of character state transition events that one or more species may have followed, which can be represented at different levels of resolution. At the highest resolution, a trajectory is a *scenario* that specifies the number, order, and timing of all character-state transitions along a lineage (Fig.1B-C). At a lower resolution, a trajectory may be represented only by its *transition sequence*, which preserves the order of state transitions, but discards their exact timing (Fig.1D-E).

**Table 1:** Definitions of key concepts and summary statistics used in this manuscript.

| Term | Definition |
| --- | --- |
| **Concepts and Structures** |  |
| Phylogenetic Character History | A mapping of character states along the nodes and branches of a phylogenetic tree. In this study, all phylogenetic character histories are derived from stochastic character mapping, but typical node-wise ancestral state reconstructions can also be used for our trajectory-wise approaches by extrapolating node states to tree edges. |
| Evolutionary Trajectory | The evolving character history for a single taxon, typically measured from the root of the phylogeny to the tip. |
| Transition Sequence | An evolutionary trajectory that only records the ordered series of character state transitions. Length is measured as the number of transitions. |
| Scenario | An evolutionary trajectory that records both the sequence and timing of character state transitions. Length is measured in phylogenetic branch length (time units). |
| Trajectory Tree | A tree of unique evolutionary trajectories. Branching reflects divergence in character histories rather than cladogenesis. Constructed by compressing a phylogenetic character history, collapsing branches that follow identical trajectories. |
| Transition Tree | A trajectory tree representing unique transition sequences, with branch lengths measured in numbers of state transitions. |
| Scenario Tree | A trajectory tree representing unique scenarios, with branch lengths measured in the same units as the phylogeny (time). |
| **Summary Statistics** |  |
| State Transition | The total number of character state transitions present in a single phylogenetic character history. |
| Lineage-Differentiating Transitions | The number of transitions where the phylogenetic lineages sharing an identical trajectory diverge due to one or more transitions to a different state. These events generate branching in the scenario tree. |
| Path-Unique Transitions | The number of transitions that follow a novel transition sequence. Equivalent to the total branch length of the transition tree, excluding the root branch. |
| Unique Transition Sequences | The number of distinct transition sequences in a phylogenetic character history. Equivalent to the number of terminal tips in the transition tree. |
| Scenario Tree Length | The total evolutionary time spent in unique scenarios. Equivalent to the total branch length of the scenario tree. |

Trajectories summarize information in phylogenetic character histories, which report state information along branches and at nodes of the phylogeny, as can be obtained from stochastic character mapping (Huelsenbeck et al. 2003; Bollback 2006). Node-wise state estimates from parsimony- and likelihood-based ancestral state reconstructions can also be used to define trajectories, provided that the states at nodes can be extrapolated as transitions along branches between nodes. Many existing transition-tree methods are based on node-wise reconstructions of this form (Ishikawa et al. 2019; Chevenet et al. 2024).

A *trajectory tree* is a tree-like representation whose tips correspond to distinct evolutionary trajectories and whose internal structure summarizes how those trajectories diverge. Trajectory trees do not introduce information beyond what is present in the underlying phylogenetic character history; rather, they reorganize that information to make patterns of character evolution more explicit and easier to summarize. When trajectories are defined as fully time-resolved scenarios, the resulting structure is a *scenario tree*, denoted Ψ*_s_* (Fig.1C). When trajectories are defined only by their transition sequences, the resulting structure is a *transition tree*, denoted Ψ*_t_* (Fig.1E).

Together, scenario trees and transition trees provide complementary compressed representations of phylogenetic character histories. Scenario trees preserve temporal information but remove explicit lineage diversification, representing diversification of scenarios rather than species. Transition trees apply a further level of compression, discarding temporal information to emphasize patterns in transition sequences alone.

Scenario trees and transition trees also differ in several respects relevant to interpretation. Scenario-tree branches are measured in units of absolute or relative time, whereas transition-tree branches are measured in numbers of transitions. Only scenario trees preserve the timing of divergence between trajectories: lineages only separate into distinct scenarios once they differ in character state at a particular time. As a result, multiple phylogenetic lineages may remain within a single scenario following speciation events if they retain the same state, whereas a lineage undergoing a state transition separates into a new scenario at the moment of that transition. Branching in a scenario tree, therefore, reflects divergence in character history rather than taxon origination, and in most cases is analogous to a budding process in state space.

Both scenario trees and transition trees may store additional information along branches beyond the current trajectory state. In our framework, each branch carries a *size*, recording the number of lineages or scenarios that contribute to it. In scenario trees, size reflects the number of phylogenetic lineages that accumulate within a specific scenario through time, and branches are labeled as *root*, *stay*, or *leave* depending on whether they originate at the root, retain the parent branch state, or represent divergence into a new state. In transition trees, size records how many phylogenetic lineages or scenarios contribute to a given transition sequence.

### Trajectory Tree Construction

Compressing a phylogenetic character history into a trajectory tree requires merging phylogenetic lineages that occupy the same discrete character state along a specific character-state transition sequence (character state transition sequence) or scenario (character state transition timings and sequence). In our approach, each phylogenetic character history (Fig. S1A) is represented by the matrix *X*, where row *i* is species *i*’s root-to-tip scenario and column *t* contains the states for each species a time *τ_t_*. Times (since the present) are represented by the ordered vector, (*τ*_0_*, τ*_1_*, . . ., τ_T_*_max_), such that *τ_t_ > τ_t_*_+1_ for all time-indexes *t*. The times in *τ* correspond to every speciation event and every character state transition in the phylogenetic character history, and the end of the process at time *T*_max_ = 0. Just as the state is recorded for every root-tip scenario, the phylogenetic edge is recorded in the phylogenetic edge matrix *E_p_*, which records the phylogenetic-tree edge index leading to tip *i* during the same interval. Together, these matrices encode, for all species, the sequence of states they traverse, the timing of all transitions, and the mapping from time intervals to phylogenetic edges (Fig. S1B,C).

A scenario tree is constructed from these matrices using a simple traversal algorithm. At the initial time *τ*_0_, the scenario tree consists of a single root edge of length 0 containing all species (Fig. S1B,C). At each subsequent time step in *τ*, the algorithm operates on the current set of scenario-tree edges, each associated with a vector of species indices. For a given scenario-tree edge, *e_si_* (where *s* indicates the edge belongs to a scenario tree, and *i* is the edge index), with associated species set, *N_l_* (*l* indicating the number correspond to the tips of phylogenetic tree, not scenario tree), the set of states for all extant species at the next time step *t* + 1 determines whether the edge persists or splits. If all species in *N_l_* share the same state, then they share the same scenario, and the corresponding edge *e_si_* is extended from time *t* to *t* + 1. If multiple states are present in *N_l_*, the edge *e_si_* is split into child edges, one for each distinct state, with each child inheriting the appropriate subset of species and phylogenetic edges from *E_p_*.

Scenario-tree splits therefore occur when lineages following the same scenario undergo divergence in state (Fig. S1D-K). Typically, this produces two daughter edges: one retaining the original state with reduced size (excluding the lineage that is now following a new scenario) and one corresponding to the newly transitioned lineage. More than two daughter edges may arise when multiple lineages undergo simultaneous state changes or when cladogenetic events induce state changes in multiple daughter lineages, as in some cladogenetic state-depedent species and extinction (ClaSSE) models (Goldberg and Igíc 2012). At least one phylogenetic speciation event must occur before a scenario-tree edge can undergo a further split. If a scenario includes only a single species and that species undergoes a state transition, a scenario tree split will not occur, as no species retain the prior state. This pattern may be common, especially in phylogenies with longer branches, and can indicate species with exceptionally unique trajectories, or instances where some trajectories were unrecoverable due to extinction or incomplete sampling of extant species.

The size and topology of scenario trees are jointly determined by speciation and state-transition rates. As the number of scenario-tree edges approaches the number of phylogenetic edges, the topology of a scenario tree converges to that of its corresponding phylogeny, with exact correspondence only when all speciation events are accompanied by cladogenetic state transitions. When state transitions occur anagenetically and are temporally decoupled from speciation, scenario trees can differ substantially from phylogenetic trees in both topology and branching order. This time-indexed, matrix-based representation provides all information required to construct scenario trees and, subsequently, transition trees, while maintaining a clear separation between the definition of trajectory-tree structures and the procedures used to compress phylogenetic character histories.

Compressing phylogenetic character histories into transition trees closely follows the procedure for scenario tree compression (Fig. S2). However, unlike scenario trees, transition trees discard temporal information and summarize character histories as ordered transition steps. Character histories are represented by root-to-tip matrices with rows indexed by species *i* ∈ S and columns indexed by transition steps *r* ∈ {1*, . . ., R*_max_}. Transition steps are measured in the number of transitions, unlike the matrices in scenario trees, which are measured in time. The state matrix *X* records the character state of each species at each step, such that *X*[*i, r*] is the state occupied by species *i* at step *r*, with a terminal marker indicating that no further transitions occur beyond that step. In addition, two edge-index matrices can be used depending on the information one wishes to store along the branches of the phylogeny: the scenario-edge matrix *E_s_*, recording the last scenario-tree edge associated with each state step, and the phylogenetic-edge matrix *E_p_*, recording the phylogenetic-tree edge occupied at that step (Fig. S2A-D). These matrices jointly encode, for all species, the ordered sequence of states they traverse, the number of lineages that accumulate in a given transition sequence step, and the number of independent instances of any given transition sequence step arising. Another difference from scenario trees is that for transition tree “splitting events”, if some species remain in a certain certain transition sequence until the present or extinction (e.g. *A* → *B*) while others continue on to a new transition sequence step (e.g. *A* → *B* → *C*), then the species remaining in *A* → *B* will form an edge in the transition tree of length zero branching from the parent edge with the same transition sequence (Fig. S2I-L). Aside from these differences, the algorithm to construct transition trees operates similarly to the one used for constructing scenario trees (Fig. S2E-L).

### Summarizing Trajectory Trees

By representing phylogenetic character histories as trees of unique trajectories, we can summarize trajectory patterns with the same statistics used for phylogenetic trees. The tip number reflects the richness of unique trajectories: in transition trees, it measures distinct transition sequences, while in scenario trees, it accounts for both the timing and the order of transitions (Table 1). Scenario trees bifurcate only when one of multiple species sharing a conserved trajectory shifts into a derived state. As such, node and tip counts represent transitions from shared ancestral states to derived states. This excludes additional transitions that occur along already unique trajectories that include only a single species (Fig.1F). Tree length, a metric of phylogenetic diversity calculated by summing all branch lengths, quantifies the total richness of evolutionary history. In scenario trees, length corresponds to the cumulative duration of unique trajectories, unweighted by lineage number, whereas in transition trees it corresponds to the number of path-unique state changes that contribute to novel transition sequences (Fig.1F). These measures are most informative when compared to the phylogenetic tree length or to the total number of transitions. For example, the ratio of scenario tree length to phylogenetic tree length indicates the fraction of evolutionary history that is unique; the ratio of scenario tree node count (lineage-differentiating transition count) relative to total state transition count reflects the proportion of state transitions that result in diverging scenarios; and the ratio of transition tree length to state transition count captures redundancy, with low ratios indicating repeated use of the same sequences and ratios near one indicating that most transitions follow unique sequences. As we demonstrate later, these statistics can be summarized tree-wide or over time, as in lineage-through-time or counts-through-time plots, or in distributions of statistics among subclades.

Previous research involving transition trees primarily generated trees from single maximum-likelihood ancestral state reconstructions at each node. We adopt an similar strategy, using the maximum *a posteriori* (MAP) character history for a distribution of stochastic character maps, individually (marginally) for each point along each branch of the phylogeny (Freyman and Höhna 2019). While useful, such reconstructions are prone to generating maps with unrealistic transition patterns, including potentially more transitions than expected in any individual map due to flickering between two equally likely states, where the “best” state at any point may differ (Freyman and Höhna 2019). An appealing alternative would be to generate a trajectory tree directly from a distribution of character maps, where any given trajectory tree branch represents a trajectory that exists in one or more of the distribution samples. For scenario trees, where branches are in units of time, this is not straightforward, as virtually zero scenarios across character history maps will follow the same exact timing of transitions, thus most scenarios will diverge early, resulting in a structure resembling more of a “bush” or “star” than a tree, with all scenarios diverging early at the first state transition, and not diverging again. For transition trees, where branches are in units of state transitions, far more trajectories will be compressed together. We call this structure a *transition summary tree*. This structure plots all transition trees’ branches and presents any and all character maps in a distribution. For each tree edge, we include the mean number of lineages accumulated and the independent scenarios that lead to that step in the transition sequence. We allow users to filter branches to show only branches with a minimum posterior probability.

Beyond constructing and quantitatively summarizing trajectory trees, we introduce multiple new approaches for evaluating and summarizing trajectory-wise patterns. Tanglegrams provide an additional means of describing the degree of congruence between phylogenies and trajectory trees. While traditionally applied to pairs of phylogenies from different taxa with shared cophylogenetic histories (Hafner and Nadler 1990), they can also be applied to compare a phylogeny with its compressed trajectory tree. Because scenario and transition trees condense both paraphyletic and monophyletic groups into shared trajectories, their topologies may differ considerably from the original phylogeny (Fig. 1 A, C). When a compressed tree is more congruent with a corresponding phylogeny, trajectory divergence times will more frequently correspond to clade ages (Fig. S3). Among lineages undergoing trajectory-diverging state transitions, older lineages will transition away earlier than younger lineages. For this reason, clades that include multiple successive trajectory divergences will often show particularly congruent patterns. Conversely, when the compressed tree is less congruent to the corresponding phylogeny, trajectory divergence will less frequently correspond to clade ages (Fig. S3). Among lineages undergoing trajectory-diverging state transitions, younger lineages may be just as likely to transition before older lineages. Following the logic above, multiple successive trajectory divergences will rarely result in incongruent patterns. A less congruent character history may include an ancestral root state that is widely conserved across the tree, and trajectory divergence arises only from state transitions away from this root state. Higher congruence may suggest a closer link between trajectory divergence in a specific character and lineage diversification, a question of great interest to many biologists.

As another example, lineage-through-time plots are a common way to compare patterns of lineage accumulation across clades and lineages in specific states (Stadler 2008; Skeels 2019; Helmstetter et al. 2022). Here, we expand on these approaches to include the accumulation of novel trajectories through time, to generate scenario-through-time and transition sequence-through-time plots. Given that scenario and transition tree branching is closely linked to state transitions, we also include state transitions through time plots. This provides a means of comparing the accumulation of path-unique and lineage-differentiating transitions, relative to all evolution transitions.

### Hypothesis Testing with Trajectory Trees

Like other summary statistics generated from phylogenetic tip data and character histories, summaries from trajectory trees can be used to test whether patterns inferred from empirical data match patterns expected given the best-fit model. In this paper, we will test the adequacy of empirical character histories generated from observed tip data and a best-fit model, to simulated character histories generated from simulated tip data and the best-fit model (Brown 2014; Brown and Thomson 2018).

Across all subsequent analyses, our approach processes trajectory tree information as follows: (1) We fit a model of character evolution to tip data; (2) Given the fitted model and tip data, we generate many stochastic character maps, transforming each into a scenario tree and transition tree; (3) We calculate our key summary statistics, including traditional character state transition count, and our new trajectory-wise statistics calculated from each scenario tree and transition tree, including tip counts, node counts, and tree lengths. We average these summary statistics across all stochastic character maps; (4) We then simulate many tip datasets from the best-fit model. For each simulated tip dataset, we repeated steps (3) and (4), resulting in many stochastic character maps, scenario trees, transition trees, and summary statistics; (5) We compare the summary statistic means across all empirical stochastic character maps to the distribution of mean summary statistics from the simulated tip data and test whether the empirical mean summary statistics fit within the expected interval of mean simulated summary statistics (within the 95% highest density interval of simulated statistic distributions) or outside the distribution. The empirical and simulated stochastic character maps can also be used in the above-mentioned through-time plots to test whether trajectory, lineage, and state transition accumulation is greater or less than what would be expected given the best-fit model of character evolution.

### Simulation Studies

As discussed above, scenario tree branching occurs only when more than one lineage evolves along the same conserved history and one lineage transitions to a different state. As such, scenario tree diversity is linked to both the rates of state transitions and net diversification. For instance, when transition rates are much higher than speciation rates, scenario trees will more closely resemble the corresponding phylogenetic tree, since they can only be as large as the phylogeny. It is also reasonable to expect that while most state transitions will increase with higher transition rates, lineage-differentiating transitions will approach an upper limit as scenario trees begin to more closely match the phylogeny. We also expect the relationships among summary statistics to vary with transition rates and net diversification rates. For instance, when transition rates are high relative to net diversification rates, any given scenario will likely accumulate multiple species before any one lineage transitions to a new state; as such, more state transitions will be lineage-differentiating. Likewise, higher transition rates relative to net diversification rates should result in a weaker relationship. Lineages will rapidly differentiate following speciation and still continue to accrue additional transitions, but without any lineage-differentiating, until the next speciation event occurs.

To demonstrate this relationship between speciation and transition rates in generating scenario trees, we simulated character histories under several 3-state character-independent diversification models with equal transition rates amongst states. Specifically, we assess the relationship between phylogenetic length, scenario tree length, and transition tree length. We simulated under model with a fixed birth rate, but variable state transition rates, with five different levels of ratios of transition rates relative (*Q*) to speciation (*λ*) rates (ex, 5:1, 2:1, 1:1, 1:2, 1:5). To simplify the relationship between transition rates and diversification rates, we used a pure birth process with death rates fixed to zero. We note that some of these rates are biologically unrealistic, particularly those in which we assume character-state transitions occur much more frequently than speciation rates. However, we present these model treatments to show the properties of phylogenetic character histories relative to trajectory trees.

In our analyses, transition and speciation rates are character-independent, making this equivalent to a character-independent multistate-dependent species extinction model (CID-MuSSE) with extinction rates fixed to zero. We repeated this experiment twice: once with tree size left to vary, by simulating a fixed time of 1.0 and filtering out character histories that do not include all three character states, and the second simulating a tree with only 300 tips. In the first set of simulations, we assessed the distribution of summary statistics across five relative rate treatments (*Q*:*λ* of 5:1, 2:1, 1:1, 1:2, 1:5); in the second, we assessed the relationships among summary statistics under three rate treatments (*Q*:*λ* of 2:1, 1:1, 1:2). We compared scenario tree length to phylogenetic tree length to evaluate the relationship between scenario tree growth and phylogeny growth. We compared the frequency of state transitions to transitions that resulted in new lineages (lineage-differentiating for scenarios and path unique for transition sequences) to evaluate the relative patterns in trajectory accumulation across models. We then compared the numbers of lineage-differentiating transitions and path-unique transitions to evaluate whether scenario-tree and transition-tree growth differ under different relative rates of character evolution and speciation. We also compared the mean state transitions per scenario tree edge with the mean phylogenetic edge length. This was done to determine whether phylogenies with larger average branch lengths are more likely to accrue state transitions that are not lineage-differentiating. Scenario tree edges with more than a single state transition result from scenarios in which a single lineage undergoes a state change, with no lineage differentiation.

In addition to summarizing scenario-tree properties, we assessed how well trajectory-tree summary statistics diagnose the adequacy of misspecified versus correctly specified fitted Markov models using our trajectory-tree summary-test framework. We focused on differences in the underlying state-transition process by simulating discrete characters on phylogenies generated by a character-state-independent Pure-Birth process. To perform our model adequacy tests, we followed steps 1–5 described in our tree summary statistics section. Here, we provide the precise simulation and inference procedures. For each replicate, we simulated a pure-birth tree (extinction = 0), then rescaled the tree so that the root-to-tip height equaled 1.0. “True” trait histories were then simulated under one of three Mk models: equal rates (ER); equal rates, early burst (ER EB); or equal rates, deadend (ER DE). Under the ER model, all state transitions are fixed to the same rate. Under the ER EB, the *Q*-matrix shares an identical structure as in the ER model, but prior to simulating character data, the tree undergoes an early burst branch length transformation *a* = −5.0 to stretch branches considerably towards the root of the phylogeny (Harmon et al. 2010). Under the ER DE model, the *Q*-matrix shares an identical structure as in the ER model, except 50% of the states are “dead-ends” and the transition rates out of these states are fixed to zero. Transition rates for all permitted off-diagonal entries of the rate matrix *Q* were sampled from independent Exponential priors with rate parameter *r* = 2.5. We used a uniform prior over root states when simulating histories and retained replicates only when all states were realized at least once, ensuring that subsequent comparisons spanned the full state space. For each generating model, we simulated 100 “true” datasets and fit all three Markov models (ER, ER EB, and ER DE) to each. For each fitted model, we generated 100 stochastic character maps given the “true” tip states. We then converted them into trajectory-tree representations and computed mean trajectory-wise and transition-wise summary statistics across all stochastic character maps. These null distributions were then compared to the empirical statistics from the true simulated histories using null distribution checks. We repeated this experiment with phylogenetic tip size set to 100, 300, and 500, with character state number set to either 2, 4, or 8. The results were then compiled by choosing the estimates for each best-fitting model for each dataset.

### Empirical Example: Greater Antillean Anolis Ecomorphs and Regions

We applied our trajectory tree framework to the Greater Antillean *Anolis* lizard phylogeny and to ecomorph and region-occupancy data. We used the time-calibrated maximum clade credibility tree from Poe et al. (2017). The Poe et al. dataset contains a phylogeny of 379 describe *Anolis* species, from which we extracted the phylogeny corresponding to 79 Greater Antillean anoles. Each species was scored as one of six ecomorph states – trunk-ground (TG), trunk (Tr), trunk-crown (TC), twig (Tw), grass-bush (GB), and crown-giant (CG) – and one of four region states – Hispaniola (Hisp), Cuba, Puerto Rico (PR), and Jamaica (Jam) – as recorded in the phytools R package (Revell 2012, 2024) and previous studies (Mahler et al. 2010). For each dataset (ecomorph and region), we fit four different continuous-time Markov Chain models using the fitMK function from phytools, including an “equal rates” (ER) model, a “symmetrical rates” (SYM) model, and an “all rates different” (ARD) model using the phytools fit.mk function. We compared models using Akaike Information Criteria (AIC; Akaike 1973). In initial analyses, AIC strongly supported the ER model for both ecomorph and region data. Under the best-fit model for ecomorphs and regions, we also simulated 500 stochastic character maps. These 500 stochastic character maps were joint character histories including region and ecomorph data, and were simulated by lumping the *Q*-matrices from our ecomorph and region data models, and restricting all “dual transitions” involving simultaneous changes in region and ecomorph state to have zero rates. These joint character histories were then subset into region- or ecomorph-specific histories. For all downstream analyses, including summary statistics and through-time plots generated from character histories, are performed both on the “joint” character histories and the character-specific histories.

We tested the adequacy of our best-fit empirical model using a similar approach to our model-misspecification tests described in the section directly above. We transformed each stochastic character map into a scenario and transition tree and calculated average summary statistics across all character maps for each model. We compared our empirical summary statistics from each model to simulated summary statistics from 500 stochastic mappings on each of 500 tip datasets simulated from each model, and test whether, for each model, the empirical summary statistic mean falls within the 95% density of the distribution of mean summary statistics from the 500 simulated datasets. We performed our summary statistics tests on both the joint character histories, including both region and ecomorph data, and character histories subsetted to include only region or ecomorph data.

To measure patterns of phylogenetic tree, scenario tree, and transition tree lineage and trajectory accumulation dynamics, we generated through-time plots for each. We generated these plots for both tree-wide accumulation and region- and ecomorph-specific patterns of accumulation. To summarize, through-time patterns from our distribution of stochastic character maps, we plotted the median through-time accumulation pattern and the upper and lower bounds of the 95% density of tip accumulation through time. To assess the relative through-time patterns from our empirical data to the expected empirical pattern, we compared the median, 95% upper and lower through-time patterns calculated from our empirical data and another set of median, 95% upper and lower through-time patterns calculated from 100 stochastic character maps from each of 100 simulated tip datasets.

For each model, we also generated a MAP character map from our empirical stochastic character maps and calculated the marginal state at each point in time and branch. Using the marginally most probable states for the phylogeny can imply unrealistic transition events (Freyman and Höhna 2019). When reviewing our MAP character maps, we did not identify any signals of obviously unrealistic events not found in any stochastic character map, including reversals to trunk-ground ecomorphs or rapid transitions between states along single branches. We transformed the MAP character map into a scenario tree and transition tree. Given transition trees are in units of character state transitions, they are readily summarizeable across many stochastic character maps, thus we made a consensus posterior summary transition tree, which includes all transition sequences with posterior probability of 10% or higher. Tanglegrams were generated between the MAP character map, scenario tree, and transition tree by constructing an association matrix that links the tips of the phylogenetic character history to the corresponding tips on the scenario and transition trees. The tanglegram was generated from the trees and association matrices using the “simmap cophylo” function from the R package phytools.

## Results

### Simulation Experiment Results

In our pure birth and character-independent MuSSE simulations, summary statistics related to evolutionary trajectories generally increases with greater transition-to-speciation rate ratios, while phylogenetic tree length did not (Fig. 2A-F). Scenario tree length (Fig. 2B) and all transition frequency statistics exhibit a modest increase in means but a much greater increase in variance with larger upper tails and larger ratios (Fig. 2D-F). Scenario-tree length (Fig. 2B) and scenario-tree internal node count (Fig. 2E) only show a clear increase in mean and variance relative to the rate-ratio until transition rates and speciation rates are one-to-one, beyond which they level off. The ratio between scenario and phylogenetic tree lengths (Fig. 2C) increases only in mean with increasing rate ratios, while the variance decreases, with the tree length ratio leveling off to roughly 0.9 at the rate ratio of five-to-one. All statistics, excluding the scenario-phylogenetic tree length ratio, are larger at the higher speciation rate treatments. Scenario to phylogenetic tree length ratios modestly decrease in the higher speciation rate treatments at low rate ratios, but the difference decreases as the rate ratio increases.

**Figure 2:**
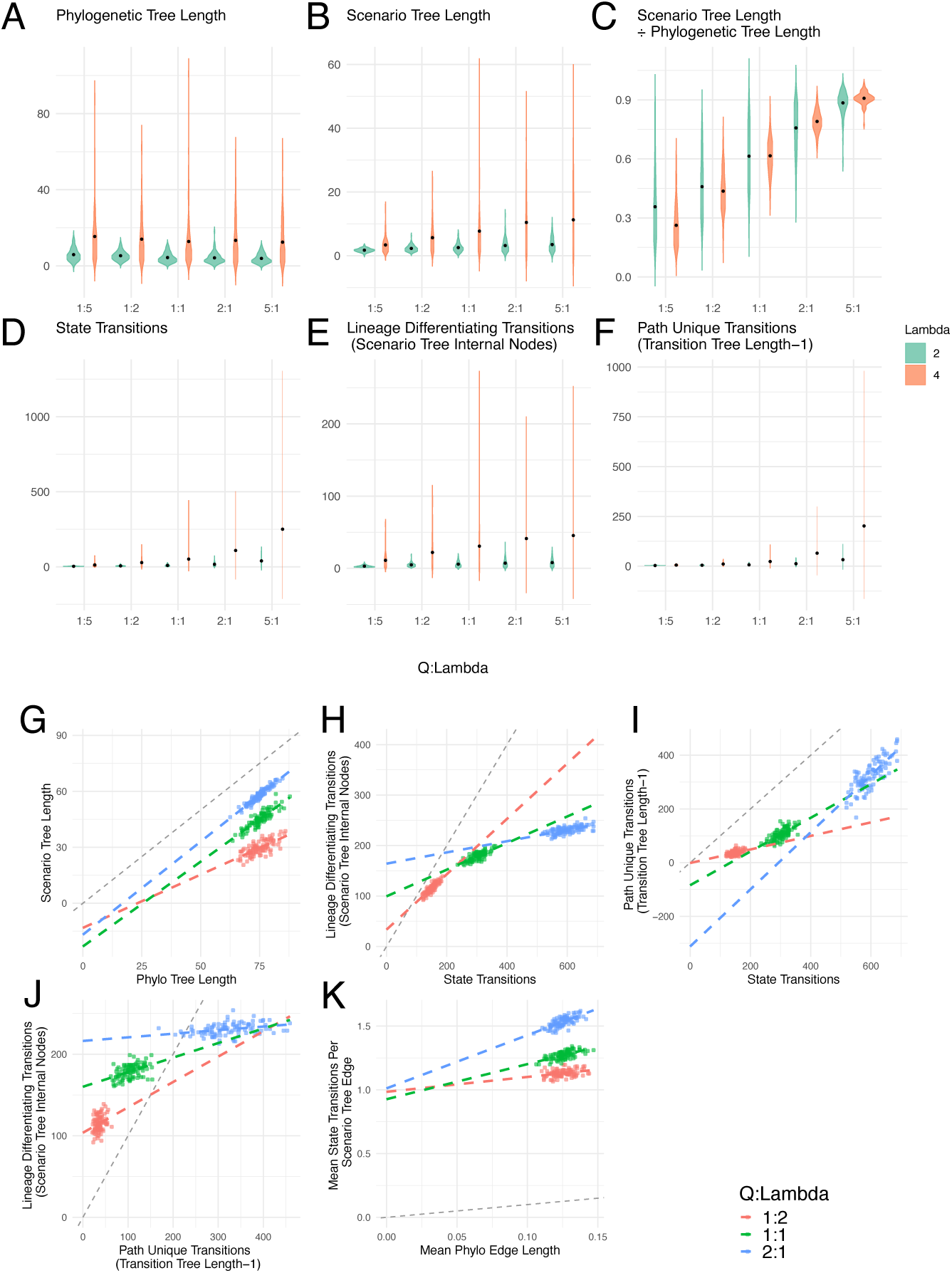
Summary statistics from three-state character histories simulated in diversitree using a Character-Independent Multistate-Dependent Speciation and Extinction Model (CID-MuSSE) In (A-F), trees in this dataset have random numbers of tips, and were filtered to only include those with character histories that involved all three states. In (G-K), trees are fixed to 300 tips. (A-F) Summary statistic distributions from character histories with speciation rate of either 2.0 or 4.0 and extinction fixed to 0. Transition rates are a ratio of the speciation rate, ranging from 1:5 to 5:1. (G-K) plots comparing relationships between summary statistics.

All summary statistics we compare show a positive correlation in our fixed-tip phylogenetic simulations, but the relationships among slopes, intercepts, and rate-ratios differ across comparisons (Fig. 2G-K). The scenario tree length versus phylogenetic tree length slope values remain roughly the same across rate-ratio treatments, while intercepts increase with increasing rate-ratios (Fig. 2G). Lineage-differentiating versus path-unique transitions, and lineage-differentiating versus all-state transitions, decrease in slope and increase in intercept as rate-ratios increase (Fig. 2H,J). The pattern reverses between path-unique state transitions and all state transitions, with slope increasing and intercept decreasing at greater rate-ratios (Fig. 2I). Also, there is a greater average number of transitions per scenario tree edge with longer phylogenetic edge lengths (Fig. 2K). This pattern becomes more prevalent as transition rates increase.

Our model misspecification test indicates that most summary statistics can discern inadequate models to some degree, often with very low rates of false positives (*<*5%) (Fig. 3). False negative rates decrease with increasing numbers of tips and state space sizes. Lineage-differentiating and path-unique transitions exhibit performance similar to that of all state transitions, but the relative number of transitions performed better, typically: path-unique transitions relative to state transitions, and lineage-differentiating transitions relative to path-unique transitions. Lineage-differentiating transitions have little power to discern inadequate from adequate models, even in our treatments, including the largest state space (8 states) and phylogenies (500 tips).

**Figure 3:**
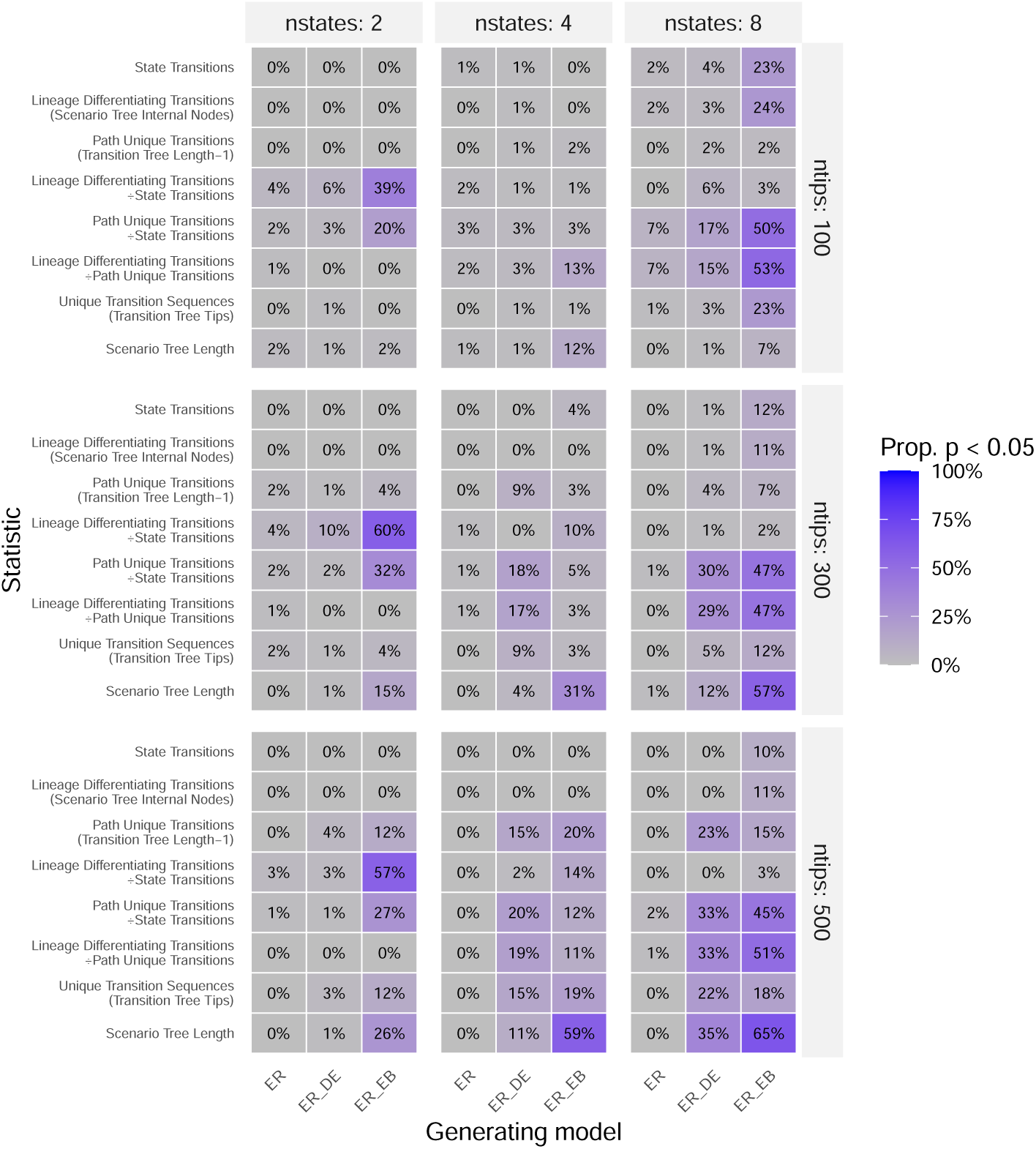
Model adequacy simulation experiment results. Tip data and character histories were simulated under 2-state, 4-state, and 8-state continuous time Markov models with either equal transition rates between all states (ER), equal rates with an “early burst” of higher rates that decrease through time (ER EB), or equal rates where some species enter, and never exit, a deadend state (ER DE). Each character history was transformed into a set of summary statistics, including our new trajectory-wise summary statistics. Each tip dataset was then fit with an ER model, and then 100 stochastic character maps were sampled given the model fit. We then compared the true empirical summary statistics from the initially simulated character histories to the distribution of summary statistics from the stochastic character maps. Higher percentages mean that stochastic character maps were less frequently able to recover the true summary statistic (ie the true statistics fall outside the 95% range of stochastic character map summary statistics generated given the ER model fit).

### Greater Antillean Anolis Region and Ecomorph Evolution

Across Greater Antillean *Anolis* lizards, we identified 9 unique biogeographic scenarios and 17 unique ecomorphological scenarios in our MAP trees (Fig. 4B, E). Region trajectories are primarily driven by the root state of Hispanola, with most other regional histories branching from this conserved state and remaining in that region until the present (Fig. 4A-C). Based on the MAP trees, ecomorph evolution is predominantly characterized by two early-diverging scenarios: the twig anole state and an early shift into the trunk-ground ecomorph (Fig. 4D-E). Both scenarios have several scenarios branching off, but the pattern is imbalanced, as the scenarios that diverge into new ecomorphs remain in those new ecomorphs until the present (Fig. 4E).

**Figure 4:**
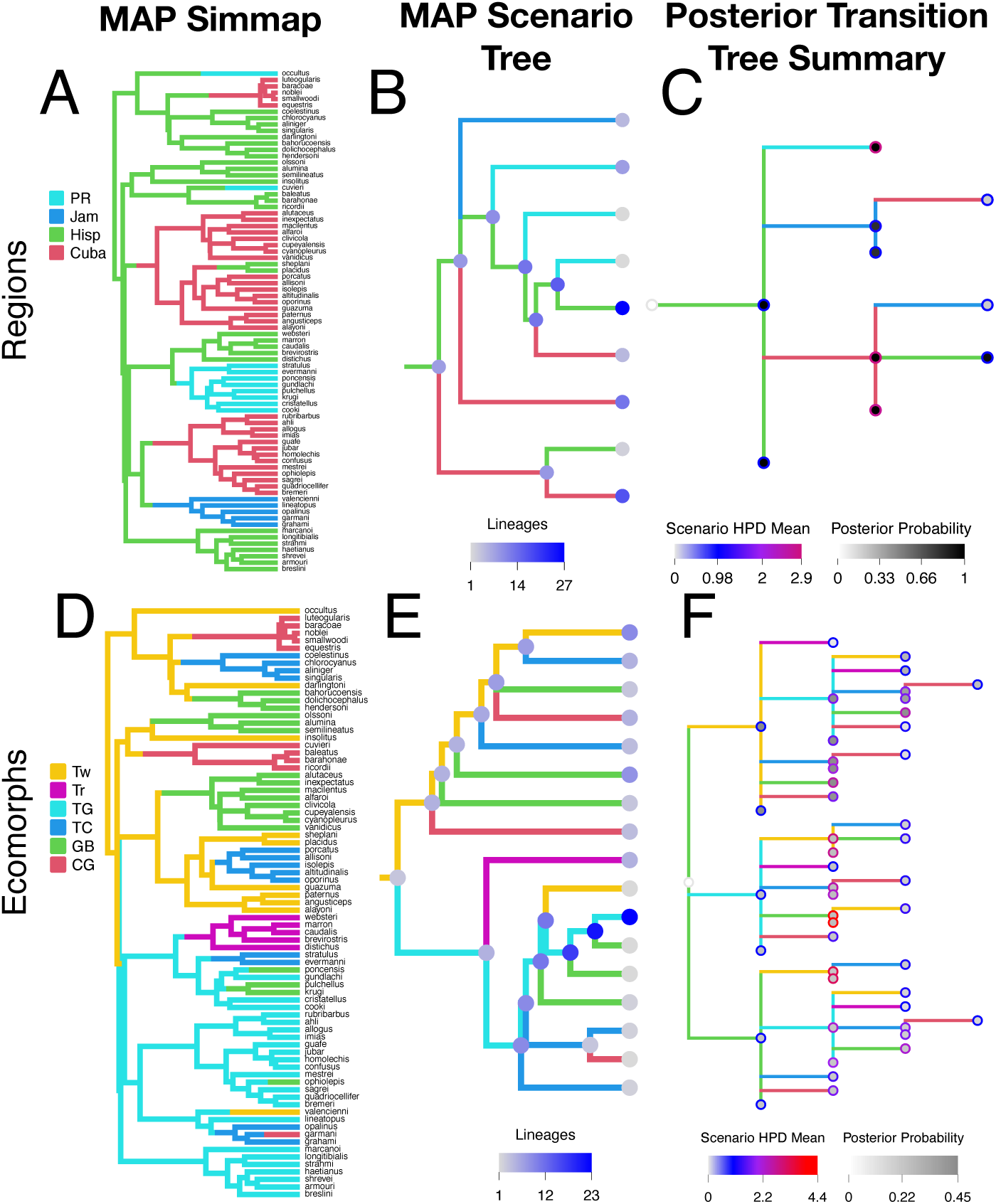
*Anolis* stochastic character maps (left), scenario trees (middle), and transition trees (right) for ecomorph and regions. Stochastic character maps and Scenario trees are both marginal *maximum a posteriori* reconstructions using the most frequently sampled state at each point along each branch of the tree. Transition trees summarize transition sequences across the distribution of 100 stochastic character maps. Node markers show the mean number of lineages (inner) that passed through at given transition sequence step, and mean number of independent evolutionary scenarios which followed a given transition sequence. Tip markers indicate the number of scenarios that terminate at a given transition sequence step.

The MAP scenario trees identified relatively unbalanced trajectories, with few transitions once a primary trajectory diverged (Fig. 4). Our transition tree posterior distribution summaries identified other subsequent transitions, albeit with lower probability (*p <* 0.5) (Fig. 4C, F). For ecomorph evolution, we identified multiple “clades” of transition sequences depending on the root state supported (Fig. 4F). In our analyses, a twig ecomorph root state had the greatest support (mean scenarios = 0.45), followed by a trunk-ground root state (mean scenarios = 0.253), with some support for a grass-bush root state (mean scenarios = 0.228). The number of transition sequences in each root-state “clade” roughly corresponded to the support for that root state (Fig. 4F). Root states with higher posterior probability had more terminal transitions with at least one independent scenario (twig:6, trunk-ground:3, grass-bush:1).

Through-time distributions summarized from showed a consistent early-burst pattern across transitions, scenarios, and transition sequence accumulation relative to simulated histories (Fig. 5 and Supplemental Figs **??**-**??**). Transition sequence through time plots showed a modestly stronger early burst pattern, with scenario accumulation noticeably greater than simulated patterns, even when normalized by tree length. Transition sequence total by the present is also relatively higher than the simulated expectation, especially compared to the number of state transitions and scenarios. State-specific accumulation patterns, including those for Cuba, grass-bush, trunk-crown, and trunk-ground lineages, are consistently higher than the mean simulated pattern (Fig. **??**-**??**). However, the through-time patterns found among other marginal state-wise through-time reconstructions were as large relative to simulated expectations as the aggregate across all states.

**Figure 5:**
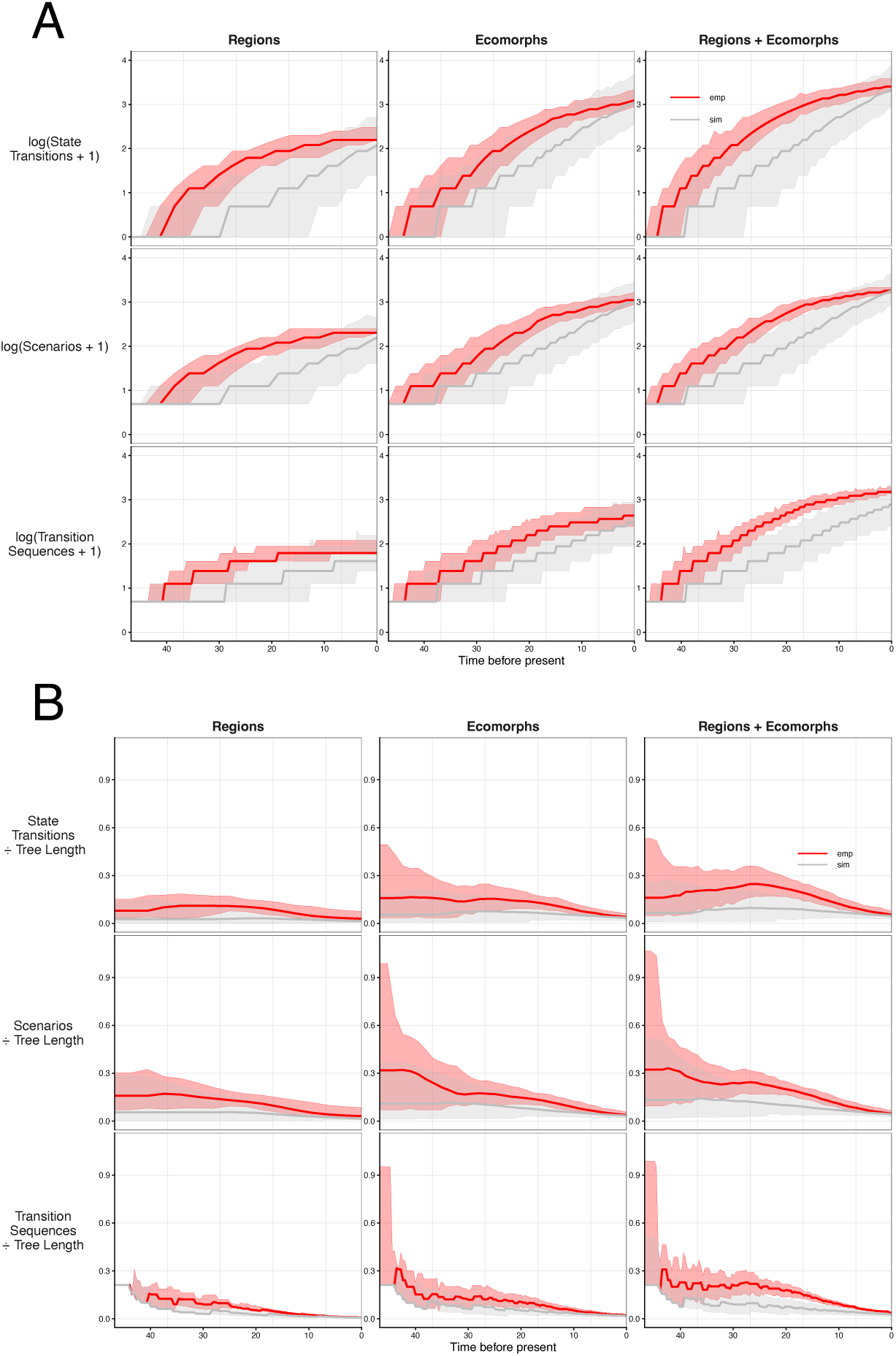
*Anolis* through-time distribution plots from the combined region and ecomorph statspace stochastic character maps for lineages, states transition, scenarios, and transition sequences. Trans-parent colors show 95% intervals while solid lines show distribution means. Each plot includes two sets of through-time distributions: an empirical distribution of stochastic character maps given the best-fit model and observed tip states (black and colored lines), and a simulated distribution (grey). Simulated distributions were generated by sampling 100 replicates of tip data from the best-fit model and the *Anolis* phylogeny, then refitting the simulated data to the best-fit model, yielding a distribution of 100 stochastic mappings for each simulated dataset. Columns display either the “full” number or the number for the indicated region or ecomorph. For region- and ecomorph-specific columns, state transitions count the number of transitions into states that in-clude the specified region or ecomorph.

**Figure 6:**
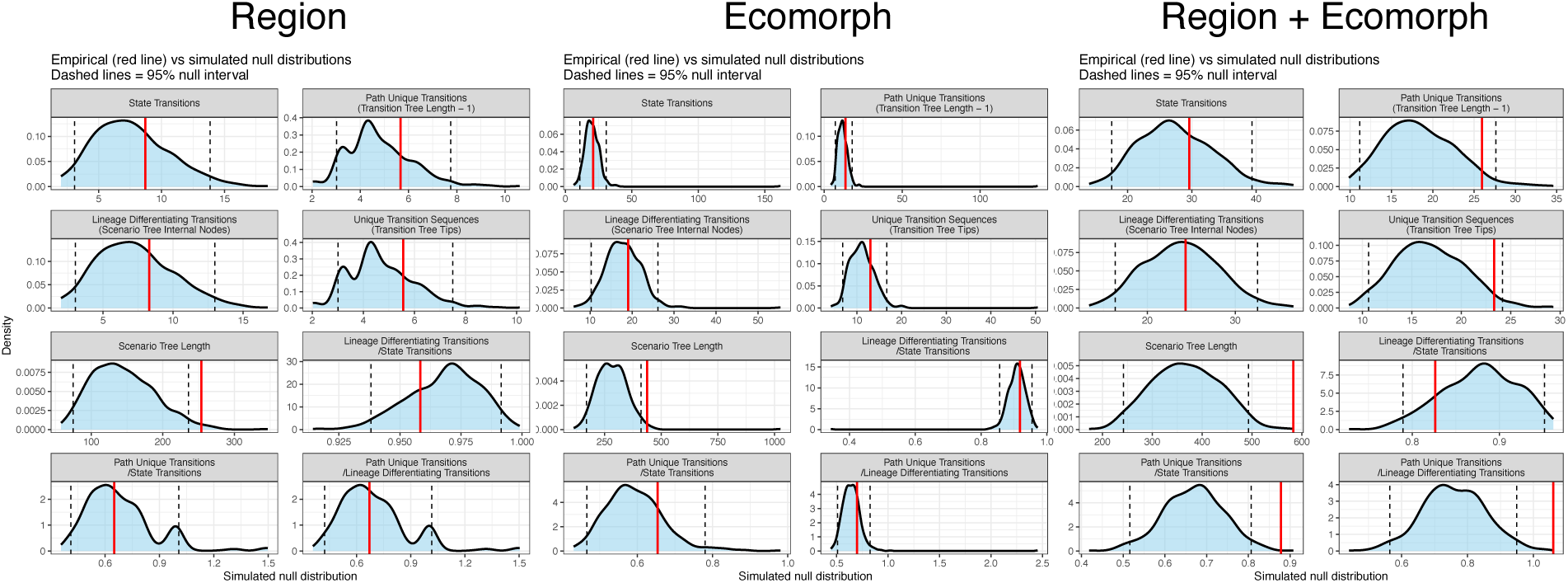
*Anolis* Simulation results, comparing mean summary statistics from a distribution of empirical stochastic character maps to a distribution of 500 mean summary statistics. Each mean summary statistic summarizes a distribution of 500 stochastic character maps inferred from fitting the best model on tip data simulated from the same model (see Methods).

Our model adequacy analysis identified only a few summary statistics that significantly deviated from simulated expectations for region- or ecomorph-specific maps (Fig. 5). One exception was the mean scenario tree length, which was significantly higher for both characters maps. For the joint region and ecomorph character maps, we found multiple mean summary statistics to be significantly higher than expected, including scenario tree length, path-unique transitions relative to the total number of state transitions, and path-unique transitions relative to the total number of lineage-differentiating transitions. While not significant, the numbers of path-unique transitions and unique transition sequences were considerably larger than the majority of simulated replicates.

Our MAP phylogeny and trajectory tree tanglegrams identified few perfectly congruent groups, but some tree pairs have far more tangles than others (Fig. 7). For region histories (Fig. 7A-B), phylogenies were largely congruent with their corresponding scenario and transition trees, but more so for scenario trees. Between the region phylogenetic character history and scenario tree, most groups were closely congruent aside from the Jamaican anoles, and the two single taxa dispersal events to Puerto Rico (*A. occultus* and *A. cuvieri*). Between the MAP region phylogenetic character history and the MAP transition tree, the Cuban crown giant anoles, and the same two single taxa dispersal to Puerto Rico form a distinct tangle. The ecomorph MAP phylogenetic character history was considerably more incongruent with either the MAP scenario or the transition tree (Fig. 7C-D). The combined region and econorph character history and MAP scenario tree is roughly intermediate in congruence between the two character histories in isolation (Fig. 7E-F).

**Figure 7:**
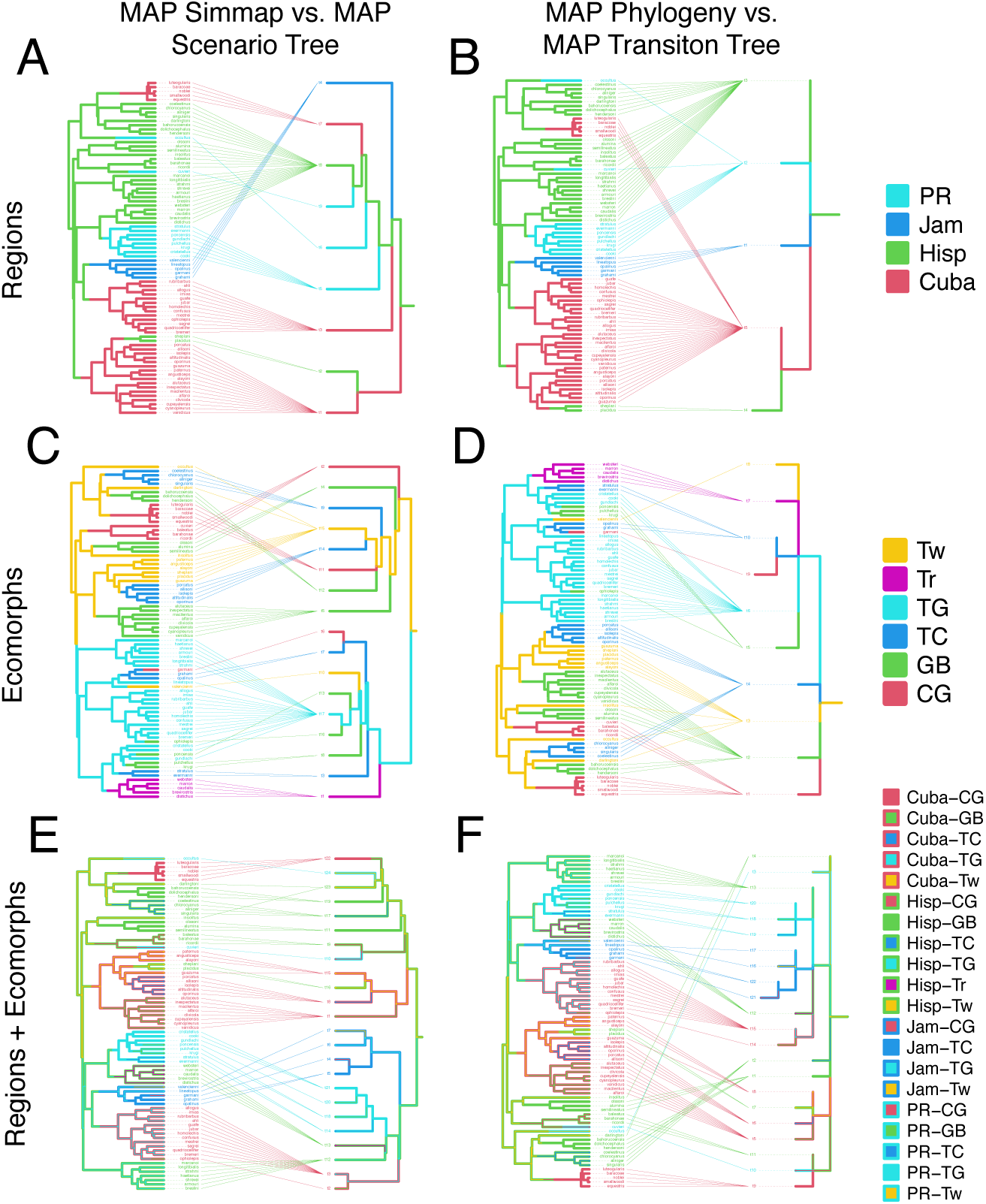
Tanglegrams between marginal maximum *a posteriori* stochastic character maps, and respective scenario trees (left) and transition trees (right) for regions (top), ecomorphs (middle), and a combined region and ecomorph state space (bottom).

The combined MAP character history is considerably more congruent for taxa branching from the twig anole root-state scenario, with the only notable tangles between *A. occultus* and the Cuban crown giant anoles, and between *A. insolitus* and two clades of Hispaniolan grass-bush (clade including *A. hendersoni*) and trunk-crown anoles (clade including *A. chlorocyanus*) (Fig. 7E-F). The major twig-to-trunk-ground scenario includes more tangles, including between the Jamaican anoles and the Hispaniolan trunk anoles and Puerto Rican anoles, and between the Cuban anole and the Hispaniolan trunk-ground anoles. The combined MAP character history and MAP transition tree are considerably incongruent with tangles across most taxa-scenario relationships.

## Discussion

Here, we introduce a framework for visually and quantitatively summarizing character transitions on phylogenies using trajectory trees: compressed phylogenetic character histories whose branches represent unique evolutionary paths. Previous work has typically used trajectory trees to represent state transition sequences, with branches in units of character-state transitions, for visualization (Chevenet et al. 2019, 2024; Ishikawa et al. 2019). To consider both the sequence and timing of character evolution, we define scenario trees and transition trees, for which lineages sharing a conserved character state history are compressed into a single lineage, also referred to as a scenario or phylotype (Chevenet et al. 2013). We also show how the branch lengths and topologies of trajectory trees, much like phylogenetic trees, have numerous, useful conceptual and quantitative properties. We measure basic tree statistics, including tree length and node numbers, to summarize the diversity of unique trajectories within a character map. We apply classic phylogenetic tree summary statistics to trajectory trees and link these statistics to specific features of character history. These include the total amount of non-conserved evolution across a tree (scenario tree length), the number of state transitions that differentiate lineages with conserved histories (scenario tree nodes − 1), the number of unique state transition sequences (transition tree tips), and the number of unique transitions within state sequences (transition tree length − 1). Using our expanded set of trajectory trees and quantitative summaries, we introduce several new approaches for identifying trajectory-wise patterns and testing whether they are within expected ranges given rates of character evolution. First, to assess whether divergence in character-evolution trajectories mirrors divergence among lineages, we use tanglegrams, a common approach for summarizing cophylogenetic patterns between clades (Matsen et al. 2018). Second, to compare how evolutionary change is distributed across lineages, states, and trajectories through time, we extend lineage- and change-through-time plots to include scenarios and transition sequences. This allows direct comparison among lineage accumulation, state accumulation, and the accumulation of unique evolutionary trajectories. Third, and lastly, to evaluate whether standard models of character evolution adequately explain the observed structure of evolutionary trajectories, we compare quantitative summaries from empirical stochastic character mappings to those obtained from simulated stochastic character mappings.

Our simulation studies identify how rates of character evolution and speciation differentially shape the diversity and complexity of trajectory trees. Scenario trees grow through lineage differentiation, which requires character-state conservatism among two or more lineages prior to state transitions. If speciation rates are too low, regardless of how fast state transitions occur, transitions would more likely occur in lineages that have already undergone at least one state transition, resulting in longer branches with multiple successive transitions and a scenario tree of similar length to a phylogeny. If speciation is much faster than state transitions, trajectories may differentiate, but given that relatively many lineages will diversify with any given trajectory, the scenario tree length will be much lower than the phylogenetic tree length.

While scenario trees are closely linked to character evolution and speciation rates, transition trees are more weakly linked to these factors. The number of transition tree tips is restricted to the number of phylogenetic tips; however, the length of transition trees is not. The total number of path-unique transitions closely matches the total number of state transitions. Regardless of whether lineages differentiate, as more successive state transitions accrue, they will build on a lineage’s existing transition sequence. When we compare path-unique transitions, we find that as transition-speciation ratios increase, a stronger correlation with total state transitions emerges. While the diversity of independent transition sequences will not increase with the number of path unique transitions, the length of these sequences will still grow, increasing the complexity of the transition tree history.

Distinguishing between the complexity and diversity of evolutionary trajectories can be useful for understanding the nature of evolutionary constraints on character evolution. For instance, when comparing transition trees between clades, what is the relationship between the number of transition tree tips and the total transition tree length? Put another way: how many transition sequences are present relative to the number of path-unique transitions present? Are there many short transition sequences, or a few long transition sequences with many steps? Do these transitions sequences diverge early or do many transition sequences share many steps and then differ on the last one or two steps? Such questions are important for understanding how constrained character evolution may be, regardless of whether rates of character evolution are fast or slow. Biologists are often want to identify the numbers and types of intermediary traits required to gain certain significant traits, and the degree of constraint acting on the order in which these intermediary steps may be followed (Donoghue 2005; Sanderson and Donoghue 1989; Donoghue and Sanderson 2015; Marazzi et al. 2012; Landis et al. 2021). If these intermediary traits are not binary and lack an obvious root state or clear pattern of accumulation (Moen and Johnston 2023; Johnston and Diaz-Uriarte 2024; Giannakis et al. 2024; Johnston et al. 2026), or are closely linked to species diversification (Maddison 2006; Maddison et al. 2007), then few other approaches exist for comparing these patterns in categorical variables.

The close relationships between trajectory patterns and rates of character evolution suggest that summary statistics from trajectory trees should have some ability to distinguish adequate from inadequate character models, even for memoryless models. We generally find that this relationship holds for most statistics, with very low false-positive rates for model adequacy (*<*5%). Importantly, the power of trajectory-wise statistics to identify inadequate models increased with the number of taxa and the size of the state space. Most statistics frequently had lower false-negative rates when fit to an equal-rates model than when fit to the dead-end model, indicating that these statistics may be more sensitive to temporal patterns in rate heterogeneity than to differences in transition-matrix structure. Our work builds on many recent important advances in evaluating model adequacy in character evolution (Schwery et al. 2023). However, most existing approaches focus only on evaluating summary statistics of tip states, while few assess the adequacy of statistics derived from character histories (Laurin-Lemay and Rodrigue 2025), with none relying on trajectory information. Our analyses explore only a small subset of all trajectory-wise summary statistics of potential use. For instance, future work should explore character state and trajectory-wise statistics, such as comparing the state diversity between phylogenic character histories and trajectory trees. Another set of statistics to consider includes tree balance statistics (Lemant et al. 2022; Janzen and Etienne 2024; Kersting et al. 2024), which can measure the balance or imbalance of trajectories. Such statistics could be used to pose questions, such as: whether most branching evolutionary trajectories stem from a single “source” to then become dead ends of disparity? or, do the derived scenarios persist and continue branching? The former would result in a ladder-like trajectory tree, while the latter would be a more balanced tree.

We emphasize that while it is useful to show that trajectory-wise summary statistics are sensitive to differences in conventional character evolution models, most models are based on simple, memoryless continuous-time Markov chains, and therefore are not designed to generate particular trajectory-wise outcomes (Goldberg and Foo 2020). We suspect that many of the trajectory-wise patterns one might identify will be difficult to explain with available character evolution models. An important direction will be to develop models that explicitly consider rates of unique and lineage-differentiating state transitions that can better explain observed patterns, possibly through the addition of time-dependent or diversity-dependent effects on the modeled rates.

Our *Anolis* analyses show that trajectory-wise character summaries readily identify distinct patterns of unique evolution that deviate from the expected outcomes of the best-fitting models of character evolution that we considered (i.e., ER over ARD and SYM). *Anolis* lizards are a classic example of an adaptive radiation, having diversified variously into a wide range of phenotypic forms and ecological niches (Losos et al. 1998; Miller et al. 2023). Previous work has generally relied on ancestral state estimation and stochastic character mapping to characterize how anoles have repeatedly dispersed and access novel niche spaces (Velasco et al. 2020; Losos 2011; Losos et al. 2006; Stroud and Losos 2016). We used evolutionary trajectories to characterize how ancestral anoles attained their current biogeographic and ecomorphological states, and whether those trajectories were concordant *Anolis* diversification. Transition sequence-through time plots show that unique transition sequences accrue across lineages faster than expected from simulations and empirical state transitions or scenarios. This is a pattern not driven by any single uptick in one advantageous state, but in aggregate across all states, consistent with the notion that adaptive radiation requires ecological differentiation among lineages. This “early burst” pattern of path-unique evolution is present in both regions and ecomorph character histories, but the pattern is strongest in combined state spaces. This pattern suggests that, to some degree, an early burst of unique evolution is greater than what might be expected by looking at transition state counts alone. This pattern of greater path-unique transitions is strongly supported by our model adequacy tests, more so than any other state transition type. In isolation, the mean number of state transitions, lineage-differentiating transitions, and path-unique transitions all fall within their respective expected simulated ranges. This pattern is only recovered in the combined region-ecomorph state space, a large state space with many different transition types. Whether this tree-wide pattern in the relative number of path-unique transitions is connected to the early burst found in our through-time analyses is unclear, as path-unique transitions could be more prevalent across the tree in general, rather than being linked to the earliest moments in the clade’s history. However, an early burst pattern in scenario divergence is still supported by significantly higher mean scenario tree lengths across empirical stochastic character maps, despite no similar signal in scenario node number relative to the number of lineage-differentiating transitions. Since the average number of nodes and tips in the scenario tree is consistent with simulated histories, the longer tree length must result from scenarios diverging earlier than expected.

Identifying whether changes in rate evolution at large match changes in the rates of unique evolution may help identify the degree to which evolutionary constraints play a role in diversification. Early burst patterns of character evolution have frequently been attributed to adaptive radiations and ecological opportunity (Harmon et al. 2010; Puttick 2018; Rundell and Price 2009; Pontarp 2021). However, these patterns are typically identified in continuous traits, not in categorical ones, and not in terms of the “uniqueness” of evolutionary events. We believe this finding raises an important question: when we observe phylogenetic variation in rates of character evolution, how often is that linked to to distinct evolutionary trajectories of character change? For instance, say we identify a multi-state character with multiple rate regimes, with some lineages evolving quickly and others evolving slowly. Depending on the redundancy of the underlying events, the path-unique transitions and rate regimes for the clade might follow the same pattern, the opposite pattern, or be completely uncorrelated. Mismatches between rates of character evolution relative to path-unique character evolution may indicate interesting patterns in character evolution constraint. This is to say that the careful use of trajectory trees may aid our understanding for how evolutionary patterns and processes give rise to uniqueness.

Phylogeny-versus-trajectory tree tanglegrams identified variable patterns of phylogenetic-trajectory tree congruence, with it being strongest between region phylogenetic character histories and scenario trees and transition trees, and weaker across all tree comparisons for ecomorphs. The combined state-space shows some congruence between the phylogenetic character history and the scenario tree, especially in scenarios stemming from the conserved twig anole scenario. These results suggest a mild relationship and an ordered structure between speciation events and lineage-differentiating transitions. However, a major limitation to this approach is the lack of uncertainty characterization in marginal MAP trees. Reporting the maximally probable state at each time step along all branches can result in considerable differences in the number of state transitions relative to those found in the underlying distribution of stochastic character maps (Freyman and Höhna 2019). While a by-eye inspection can detect obvious signs of summary error, specifically rapid “flickering” between states, this still oversimplifies the distribution of stochastic character maps. Additionally, tanglegrams alone can be prone to misidentifying the degree of topological congruence in all cases (De Vienne 2019). A potentially promising future direction to resolve both issues would be to utilize a summary statistic for cophylogenetic signal (Balbuena et al. 2020; Llaberia-Robledillo et al. 2023).

## Conclusion

Evolutionary scenarios represent complex historical sequences of evolutionary changes that produce phylogenetic diversity. How biologists identify, describe, and evaluate support for different evolutionary scenarios is crucial to our field, as it directly impacts how we capture and communicate the essence of a complex evolutionary history, and how we compare which explanations for that history are most plausible (Greene 2017). Although biologists often reason with evolutionary trajectories in terms of individual lineages, we rarely summarize trajectory properties across multiple lineages, including the number, age, size, or topology of distinct trajectories. While much work remains to formalize the mathematical properties of evolutionary trajectories, we believe the pattern-based approaches introduced by our study demonstrate that trajectory trees, in their current form, can enrich many phylogenetic studies.

## Supplementary Information

All supplementary documents and figures are available at Data Dryad at this link: TBD, Code available at https://github.com/sean-mchugh/Scenario_tree_ms_analyses/tree/main

## Acknowledgments

We are indebted to Albert Soewongsono, Jake Berv, Sarah Swiston, Tasman Ezra, Shiyang Wu, Justin Baldwin, Raymond Castillo, Anna Nagel, Bailey Howell, and Nic Bone for their insights on how to improve our presentation of the project.

## References

Olav N. L. Aga, Morten Brun, Kazeem A. Dauda, Ramon Diaz-Uriarte, Konstantinos Giannakis, and Iain G. Johnston. HyperTraPS-CT: Inference and prediction for accumulation pathways with flexible data and model structures. PLOS Computational Biology, 20(9):e1012393, September 2024. ISSN 1553-7358. doi: 10.1371/journal.pcbi.1012393. URL https://journals.plos.org/ploscompbiol/article?id=10.1371/journal.pcbi.1012393. Publisher: Public Library of Science.

Hirotugu Akaike. Information theory and an extension of the maximum likelihood principle. In Second International Symposium on Information Theory, 1973, pages 267–281. Akademia Kiado, 1973.

J. W. Archie and J. Felsenstein. The Number of Evolutionary Steps on Random and Minimum Length Trees for Random Evolutionary Data. Theoretical Population Biology, 43(1):52–79, February 1993. ISSN 0040-5809. doi: 10.1006/tpbi.1993.1003. URL https://www.sciencedirect.com/science/article/pii/S0040580983710038.

Simon Baeckens, Charlotte Goeyers, and Raoul Van Damme. Convergent Evolution of Claw Shape in a Transcontinental Lizard Radiation. Integrative and Comparative Biology, 60(1): 10–23, July 2020. ISSN 1540-7063. doi: 10.1093/icb/icz151. URL https://doi.org/10.1093/icb/icz151.

Juan Antonio Balbuena, Óscar Alejandro Pérez-Escobar, Cristina Llopis-Belenguer, and Isabel Blasco-Costa. Random Tanglegram Partitions (Random TaPas): An Alexandrian Approach to the Cophylogenetic Gordian Knot. Systematic Biology, 69(6):1212–1230, November 2020. ISSN 1063-5157, 1076-836X. doi: 10.1093/sysbio/syaa033. URL https://academic.oup.com/sysbio/article/69/6/1212/5820982.

Jeremy M. Beaulieu and Brian C. O’Meara. Detecting Hidden Diversification Shifts in Models of Trait-Dependent Speciation and Extinction. Systematic Biology, 65(4):583–601, July 2016. ISSN 1063-5157. doi: 10.1093/sysbio/syw022. URL https://doi.org/10.1093/sysbio/syw022.

Jacob S. Berv, Sonal Singhal, Daniel J. Field, Nathanael Walker-Hale, Sean W. McHugh, J. Ryan Shipley, Eliot T. Miller, Rebecca T. Kimball, Edward L. Braun, Alex Dornburg, C. Tomomi Parins-Fukuchi, Richard O. Prum, Benjamin M. Winger, Matt Friedman, and Stephen A. Smith. Genome and life-history evolution link bird diversification to the end-Cretaceous mass extinction. Science Advances, 10(31):eadp0114, August 2024. ISSN 2375-2548. doi: 10.1126/sciadv.adp0114. URL https://www.science.org/doi/10.1126/sciadv.adp0114.

Bonnie B. Blaimer, Bernardo F. Santos, Astrid Cruaud, Michael W. Gates, Robert R. Kula, Istváan Mikáo, Jean-Yves Rasplus, David R. Smith, Elijah J. Talamas, Séan G. Brady, and Matthew L. Buffington. Key innovations and the diversification of Hymenoptera. Nature Communications, 14(1):1212, March 2023. ISSN 2041-1723. doi: 10.1038/s41467-023-36868-4. URL https://www.nature.com/articles/s41467-023-36868-4. Publisher: Nature Publishing Group.

Jonathan P. Bollback. SIMMAP: Stochastic character mapping of discrete traits on phylogenies. BMC Bioinformatics, 7(1):88, February 2006. ISSN 1471-2105. doi: 10.1186/1471-2105-7-88. URL https://doi.org/10.1186/1471-2105-7-88.

Ronald M. Bonett, Nicholus M. Ledbetter, Alexander J. Hess, Madison A. Herrboldt, and Mathieu Denöel. Repeated ecological and life cycle transitions make salamanders an ideal model for evolution and development. Developmental Dynamics, 251(6):957–972, 2022. ISSN 1097-0177. doi: 10.1002/dvdy.373. URL https://onlinelibrary.wiley.com/doi/abs/10.1002/dvdy.373. eprint: https://anatomypubs.onlinelibrary.wiley.com/doi/pdf/10.1002/dvdy.373.

James D Boyko and Jeremy M Beaulieu. Reducing the Biases in False Correlations Between Discrete Characters. Systematic Biology, 72(2):476–488, March 2023. ISSN 1063-5157. doi: 10.1093/sysbio/syac066. URL https://doi.org/10.1093/sysbio/syac066.

Jeremy M Brown. Detection of implausible phylogenetic inferences using posterior predictive assessment of model fit. Systematic biology, 63(3):334–348, 2014.

Jeremy M. Brown and Robert C. Thomson. Evaluating Model Performance in Evolutionary Biology. Annual Review of Ecology, Evolution, and Systematics, 49(1):95–114, November 2018. ISSN 1543-592X, 1545-2069. doi: 10.1146/annurev-ecolsys-110617-062249. URL https://www.annualreviews.org/doi/10.1146/annurev-ecolsys-110617-062249.

F Chevenet, D Fargette, P Bastide, T Vitŕe, and S Guindon. EvoLaps 2: Advanced phylogeographic visualization. Virus Evolution, 10(1):vead078, January 2024. ISSN 2057-1577. doi: 10.1093/ve/vead078. URL https://academic.oup.com/ve/article/doi/10.1093/ve/vead078/7479648.

François Chevenet, Matthieu Jung, Martine Peeters, Tulio de Oliveira, and Olivier Gascuel. Searching for virus phylotypes. Bioinformatics, 29(5):561–570, March 2013. ISSN 1367-4803.doi: 10.1093/bioinformatics/btt010. URL https://doi.org/10.1093/bioinformatics/btt010.

François Chevenet, Guillaume Castel, Emmanuelle Jousselin, and Olivier Gascuel. PastView: a user-friendly interface to explore ancestral scenarios. BMC Evolutionary Biology, 19(1):163, December 2019. ISSN 1471-2148. doi: 10.1186/s12862-019-1490-4. URL https://bmcevolbiol.biomedcentral.com/articles/10.1186/s12862-019-1490-4.

Kristen E. Crandell, Anthony Herrel, Mahmood Sasa, Jonathan B. Losos, and Kellar Autumn. Stick or grip? Co-evolution of adhesive toepads and claws in Anolis lizards. Zoology, 117(6): 363–369, December 2014. ISSN 0944-2006. doi: 10.1016/j.zool.2014.05.001. URL https://www.sciencedirect.com/science/article/pii/S0944200614000610.

Damien M De Vienne. Tanglegrams Are Misleading for Visual Evaluation of Tree Congruence. Molecular Biology and Evolution, 36(1):174–176, January 2019. ISSN 0737-4038, 1537-1719. doi: 10.1093/molbev/msy196. URL https://academic.oup.com/mbe/article/36/1/174/5142657.

Michael J. Donoghue. Key innovations, convergence, and success: macroevolutionary lessons from plant phylogeny. Paleobiology, 31(S2):77–93, January 2005. ISSN 0094-8373, 1938-5331. doi: 10.1666/0094-8373(2005)031[0077:KICASM]2.0.CO;2. URL https://www.cambridge.org/core/journals/paleobiology/article/abs/key-innovations-convergence-and-success-macroevolutionary-lessons-from-plant-phylogeny/2DDAFB62E85BB38E84E995E57B6A9612.

Michael J. Donoghue and Joachim W. Kadereit. Walter Zimmermann and the growth of phylogenetic theory. Systematic Biology, 41(1):74–85, 1992.

Michael J. Donoghue and Michael J. Sanderson. Confluence, synnovation, and depauperons in plant diversification. New Phytologist, 207(2):260–274, 2015. ISSN 1469-8137. doi: 10.1111/nph.13367. URL https://onlinelibrary.wiley.com/doi/abs/10.1111/nph.13367. eprint: https://onlinelibrary.wiley.com/doi/pdf/10.1111/nph.13367.

Douglas H. Erwin. Novelty and Innovation in the History of Life. Current Biology, 25(19): R930–R940, October 2015. ISSN 09609822. doi: 10.1016/j.cub.2015.08.019. URL https://linkinghub.elsevier.com/retrieve/pii/S0960982215009902.

Damien Esquerŕe and J. Scott Keogh. Parallel selective pressures drive convergent diversification of phenotypes in pythons and boas. Ecology Letters, 19(7):800–809, 2016. ISSN 1461-0248. doi: 10.1111/ele.12620. URL https://onlinelibrary.wiley.com/doi/abs/10.1111/ele.12620. eprint: https://onlinelibrary.wiley.com/doi/pdf/10.1111/ele.12620.

James S. Farris. The Retention Index and Homoplasy Excess. Systematic Zoology, 38(4):406, December 1989. ISSN 00397989. doi: 10.2307/2992406. URL https://academic.oup.com/sysbio/article-lookup/doi/10.2307/2992406.

William A Freyman and Sebastian Höhna. Stochastic Character Mapping of State-Dependent Diversification Reveals the Tempo of Evolutionary Decline in Self-Compatible Onagraceae Lineages. Systematic Biology, 68(3):505–519, May 2019. ISSN 1063-5157. doi: 10.1093/sysbio/syy078. URL https://doi.org/10.1093/sysbio/syy078.

Tony Gamble, Eli Greenbaum, Todd R. Jackman, Anthony P. Russell, and Aaron M. Bauer. Repeated origin and loss of adhesive toepads in geckos. PLoS One, 7(6):e39429, 2012.

Konstantinos Giannakis, Olav N. L. Aga, Marcus T. Moen, Pål G. Drange, and Iain G. Johnston. Identifying parsimonious pathways of accumulation and convergent evolution from binary data. bioRxiv, 2024. doi: 10.1101/2024.11.06.622201. URL https://www.biorxiv.org/content/early/2024/11/08/2024.11.06.622201.

Emma E. Goldberg and Jasmine Foo. Memory in Trait Macroevolution. The American Naturalist, 195(2):300–314, February 2020. ISSN 0003-0147. doi: 10.1086/705992. URL https://www.journals.uchicago.edu/doi/full/10.1086/705992. Publisher: The University of Chicago Press. Emma E Goldberg and Boris Igíc. Tempo and mode in plant breeding system evolution. *Evolution*, 66(12):3701–3709, 2012.

Sam F. Greenbury, Mauricio Barahona, and Iain G. Johnston. HyperTraPS: Inferring Probabilistic Patterns of Trait Acquisition in Evolutionary and Disease Progression Pathways. Cell Systems, 10(1):39–51.e10, January 2020. ISSN 2405-4712. doi: 10.1016/j.cels.2019.10.009. URL https://www.sciencedirect.com/science/article/pii/S2405471219303850.

Harry W. Greene. Evolutionary Scenarios and Primate Natural History. The American Naturalist, 190(S1):S69–S86, August 2017. ISSN 0003-0147, 1537-5323. doi: 10.1086/692830. URL https://www.journals.uchicago.edu/doi/10.1086/692830.

Aaron H Griffing, Tony Gamble, Martin J Cohn, and Thomas J Sanger. Convergent developmental patterns underlie the repeated evolution of adhesive toe pads among lizards. Biological Journal of the Linnean Society, 135(3):518–532, March 2022. ISSN 0024-4066. doi: 10.1093/biolinnean/blab164. URL https://doi.org/10.1093/biolinnean/blab164.

Mark S Hafner and Steven A Nadler. Cospeciation in host-parasite assemblages: comparative analysis of rates of evolution and timing of cospeciation events. Systematic Zoology, 39(3): 192–204, 1990.

Mattias Hagman and Terry J. Ord. Many Paths to a Common Destination: Morphological Differentiation of a Functionally Convergent Visual Signal. The American Naturalist, 188(3): 306–318, September 2016. ISSN 0003-0147, 1537-5323. doi: 10.1086/687560. URL https://www.journals.uchicago.edu/doi/10.1086/687560.

Luke J. Harmon, Jonathan B. Losos, T. Jonathan Davies, Rosemary G. Gillespie, John L. Gittleman, W. Bryan Jennings, Kenneth H. Kozak, Mark A. McPeek, Franck Moreno-Roark, Thomas J. Near, Andy Purvis, Robert E. Ricklefs, Dolph Schluter, James A. Schulte Ii, Ole Seehausen, Brian L. Sidlauskas, Omar Torres-Carvajal, Jason T. Weir, and Arne Ø Mooers. Early Bursts of Body Size and Shape Evolution Are Rare in Comparative Data. Evolution, 64 (8):2385–2396, 2010. ISSN 1558-5646. doi: 10.1111/j.1558-5646.2010.01025.x. URL https://onlinelibrary.wiley.com/doi/abs/10.1111/j.1558-5646.2010.01025.x. eprint: https://onlinelibrary.wiley.com/doi/pdf/10.1111/j.1558-5646.2010.01025.x.

Michael Hautmann. What is macroevolution? Palaeontology, 63(1):1–11, January 2020. ISSN 0031-0239, 1475-4983. doi: 10.1111/pala.12465. URL https://onlinelibrary.wiley.com/doi/10.1111/pala.12465.

Andrew J Helmstetter, Sylvain Glemin, Jos Käfer, Rosana Zenil-Ferguson, Hervé Sauquet, Hugo De Boer, Ĺeo-Paul M J Dagallier, Nathan Mazet, Eliette L Reboud, Thomas L P Couvreur, and Fabien L Condamine. Pulled Diversification Rates, Lineages-Through-Time Plots, and Modern Macroevolutionary Modeling. Systematic Biology, 71(3):758–773, April 2022. ISSN 1063-5157, 1076-836X. doi: 10.1093/sysbio/syab083. URL https://academic.oup.com/sysbio/article/71/3/758/6382322.

Jennifer F. Hoyal Cuthill. The morphological state space revisited: what do phylogenetic patterns in homoplasy tell us about the number of possible character states? Interface Focus, 5(6): 20150049, December 2015. doi: 10.1098/rsfs.2015.0049. URL https://royalsocietypublishing.org/doi/10.1098/rsfs.2015.0049. Publisher: Royal Society.

John P Huelsenbeck, Rasmus Nielsen, and Jonathan P Bollback. Stochastic Mapping of Morphological Characters. SYSTEMATIC BIOLOGY, 52, 2003.

Jonathan M Huie, Ivan Prates, Rayna C Bell, and Kevin De Queiroz. Convergent patterns of adaptive radiation between island and mainland *Anolis* lizards. Biological Journal of the Linnean Society, 134(1):85–110, August 2021. ISSN 0024-4066, 1095-8312. doi: 10.1093/biolinnean/blab072. URL https://academic.oup.com/biolinnean/article/134/1/85/6287635.

John P Hunter. Key innovations and the ecology of macroevolution. Trends in Ecology & Evolution, 13(1):31–36, January 1998. ISSN 0169-5347. doi: 10.1016/S0169-5347(97)01273-1. URL https://www.sciencedirect.com/science/article/pii/S0169534797012731.

Sohta A Ishikawa, Anna Zhukova, Wataru Iwasaki, and Olivier Gascuel. A Fast Likelihood Method to Reconstruct and Visualize Ancestral Scenarios. Molecular Biology and Evolution, 36 (9):2069–2085, September 2019. ISSN 0737-4038. doi: 10.1093/molbev/msz131. URL https://doi.org/10.1093/molbev/msz131.

Thijs Janzen and Rampal S. Etienne. Phylogenetic tree statistics: A systematic overview using the new R package ‘treestats’. Molecular Phylogenetics and Evolution, 200:108168, November 2024. ISSN 1055-7903. doi: 10.1016/j.ympev.2024.108168. URL https://www.sciencedirect.com/science/article/pii/S105579032400160X.

Iain Johnston, Ramon Diaz-Uriarte, and James Boyko. The hypercubic mk model in reduced state space for the coupled, reversible coevolution of multiple binary characters. bioRxiv, 2026. doi: 10.64898/2026.06.01.729317. URL https://www.biorxiv.org/content/early/2026/06/03/2026.06.01.729317.

Iain G Johnston and Ramon Diaz-Uriarte. A hypercubic Mk model framework for capturing reversibility in disease, cancer, and evolutionary accumulation modelling. Bioinformatics, 41 (1):btae737, December 2024. ISSN 1367-4811. doi: 10.1093/bioinformatics/btae737. URL https://academic.oup.com/bioinformatics/article/doi/10.1093/bioinformatics/btae737/7922554.

Sophie J. Kersting, Kristina Wicke, and Mareike Fischer. Tree balance in phylogenetic models, October 2024. URL http://arxiv.org/abs/2406.05185. arXiv:2406.05185 [q-bio].

Michael Landis, Erika J Edwards, and Michael J Donoghue. Modeling Phylogenetic Biome Shifts on a Planet with a Past. Systematic Biology, 70(1):86–107, January 2021. ISSN 1063-5157, 1076-836X. doi: 10.1093/sysbio/syaa045. URL https://academic.oup.com/sysbio/article/70/1/86/5854876.

Simon Laurin-Lemay and Nicolas Rodrigue. Stochastic Character Mapping: An Under-Exploited Approach to the Study of Molecular Evolution. Journal of Molecular Evolution, 93(4):465–473, 2025. ISSN 0022-2844. doi: 10.1007/s00239-025-10257-5. URL https://pmc.ncbi.nlm.nih.gov/articles/PMC12354545/.

Jeanne Lemant, Ćecile Le Sueur, Veselin Manojlovíc, and Robert Noble. Robust, Universal Tree Balance Indices. Systematic Biology, 71(5):1210–1224, September 2022. ISSN 1063-5157. doi: 10.1093/sysbio/syac027. URL https://doi.org/10.1093/sysbio/syac027.

Philippe Lemey, Vladimir N. Minin, Filip Bielejec, Sergei L. Kosakovsky Pond, and Marc A. Suchard. A counting renaissance: combining stochastic mapping and empirical Bayes to quickly detect amino acid sites under positive selection. Bioinformatics, 28(24):3248–3256, December 2012. ISSN 1367-4803. doi: 10.1093/bioinformatics/bts580. URL https://doi.org/10.1093/bioinformatics/bts580.

Paul O. Lewis. A Likelihood Approach to Estimating Phylogeny from Discrete Morphological Character Data. Systematic Biology, 50(6):913–925, November 2001. ISSN 1076-836X, 1063-5157. doi: 10.1080/106351501753462876. URL http://academic.oup.com/sysbio/article/50/6/913/1628902/A-Likelihood-Approach-to-Estimating-Phylogeny-from.

Mar Llaberia-Robledillo, J Ignacio Lucas-Lledáo, Oscar A Pérez-Escobar, Boris R Krasnov, and Juan Antonio Balbuena. Rtapas: An R Package to Assess Cophylogenetic Signal between Two Evolutionary Histories. Systematic Biology, 72(4):946–954, July 2023. ISSN 1063-5157. doi: 10.1093/sysbio/syad016. URL https://doi.org/10.1093/sysbio/syad016.

Jonathan B. Losos. Ecomorphology, Performance Capability, and Scaling of West Indian Anolis Lizards: An Evolutionary Analysis. Ecological Monographs, 60(3):369–388, September 1990. ISSN 0012-9615, 1557-7015. doi: 10.2307/1943062. URL https://esajournals.onlinelibrary.wiley.com/doi/10.2307/1943062.

Jonathan B Losos. THE EVOLUTION OF CONVERGENT STRUCTURE IN CARIBBEAN ANOLIS COMMUNITIES. Systematic Biology, 41, 1992.

Jonathan B. Losos. Lizards in an Evolutionary Tree: Ecology and Adaptive Radiation of Anoles. Univ of California Press, February 2011. ISBN 978-0-520-26984-2. Google-Books-ID: SakwDwAAQBAJ.

Jonathan B. Losos, Todd R. Jackman, Allan Larson, Kevin de Queiroz, and Lourdes Rodrıguez-Schettino. Contingency and Determinism in Replicated Adaptive Radiations of Island Lizards. Science, 279(5359):2115–2118, March 1998. doi: 10.1126/science.279.5359.2115. URL https://www.science.org/doi/full/10.1126/science.279.5359.2115. Publisher: American Association for the Advancement of Science.

Jonathan B. Losos, Richard E. Glor, Jason J. Kolbe, and Kirsten Nicholson. ADAPTATION, SPECIATION, AND CONVERGENCE: A HIERARCHICAL ANALYSIS OF ADAPTIVE RADIATION IN CARIBBEAN ANOLIS LIZARDS^1^. Annals of the Missouri Botanical Garden, 93(1):24–33, May 2006. ISSN 0026-6493. doi: 10.3417/0026-6493(2006)93[24:ASACAH]2.0.CO;2. URL http://www.bioone.org/doi/abs/10.3417/0026-6493%282006%2993%5B24%3AASACAH%5D2.0.CO%3B2.

Wayne P. Maddison. Confounding Asymmetries in Evolutionary Diversification and Character Change. Evolution, 60(8):1743–1746, 2006. ISSN 1558-5646. doi: 10.1111/j.0014-3820.2006.tb00517.x. URL https://onlinelibrary.wiley.com/doi/abs/10.1111/j.0014-3820.2006.tb00517.x. eprint: https://onlinelibrary.wiley.com/doi/pdf/10.1111/j.0014-3820.2006.tb00517.x.

Wayne P Maddison, Peter E Midford, and Sarah P Otto. Estimating a Binary Character’s Effect on Speciation and Extinction. SYSTEMATIC BIOLOGY, 56, 2007.

D. Luke Mahler, Liam J. Revell, Richard E. Glor, and Jonathan B. Losos. Ecological opportunity and the rate of morphological evolution in the diversification of Greater Antillean Anolis. Evolution, 64(9):2731–2745, September 2010. ISSN 0014-3820. doi: 10.1111/j.1558-5646.2010.01026.x. URL https://doi.org/10.1111/j.1558-5646.2010.01026.x.

Brigitte Marazzi, Ćecile Ańe, Marcelo F. Simon, Alfonso Delgado-Salinas, Melissa Luckow, and Michael J. Sanderson. Locating Evolutionary Precursors on a Phylogenetic Tree. Evolution, 66 (12):3918–3930, 2012. ISSN 1558-5646. doi: 10.1111/j.1558-5646.2012.01720.x. URL https://onlinelibrary.wiley.com/doi/abs/10.1111/j.1558-5646.2012.01720.x. eprint: https://onlinelibrary.wiley.com/doi/pdf/10.1111/j.1558-5646.2012.01720.x.

Frederick A. Matsen, Sara C. Billey, Arnold Kas, and Matja_̌_z Konvalinka. Tanglegrams: A Reduction Tool for Mathematical Phylogenetics. IEEE/ACM Transactions on Computational Biology and Bioinformatics, 15(1):343–349, January 2018. ISSN 1557-9964. doi: 10.1109/TCBB.2016.2613040. URL https://ieeexplore.ieee.org/document/7581044/?arnumber=7581044. Conference Name: IEEE/ACM Transactions on Computational Biology and Bioinformatics.

Michael R. May and Bruce Rannala. Early detection of highly transmissible viral variants using phylogenomics. Science Advances, 10(33):eadk7623, August 2024. doi: 10.1126/sciadv.adk7623. URL https://www.science.org/doi/full/10.1126/sciadv.adk7623. Publisher: American Association for the Advancement of Science.

M. F. Mickevich and S. J. Weller. Evolutionary Character Analysis: Tracing Character Change on a Cladogram. Cladistics, 6(2):137–170, 1990. ISSN 1096-0031. doi: 10.1111/j.1096-0031.1990.tb00533.x. URL https://onlinelibrary.wiley.com/doi/abs/10.1111/j.1096-0031.1990.tb00533.x. eprint: https://onlinelibrary.wiley.com/doi/pdf/10.1111/j.1096-0031.1990.tb00533.x.

Aryeh H Miller and James T Stroud. Novel Tests of the Key Innovation Hypothesis: Adhesive Toepads in Arboreal Lizards. Systematic Biology, page syab041, June 2021. ISSN 1063-5157, 1076-836X. doi: 10.1093/sysbio/syab041. URL https://academic.oup.com/sysbio/advance-article/doi/10.1093/sysbio/syab041/6295694.

Aryeh H. Miller, James T. Stroud, and Jonathan B. Losos. The ecology and evolution of key innovations. Trends in Ecology & Evolution, 38(2):122–131, February 2023. ISSN 01695347. doi: 10.1016/j.tree.2022.09.005. URL https://linkinghub.elsevier.com/retrieve/pii/S0169534722002257.

Emily Mitchell, Michael Spencer Chapman, Nicholas Williams, Kevin J. Dawson, Nicole Mende, Emily F. Calderbank, Hyunchul Jung, Thomas Mitchell, Tim H. H. Coorens, David H. Spencer, Heather Machado, Henry Lee-Six, Megan Davies, Daniel Hayler, Margarete A. Fabre, Krishnaa Mahbubani, Federico Abascal, Alex Cagan, George S. Vassiliou, Joanna Baxter, Inigo Martincorena, Michael R. Stratton, David G. Kent, Krishna Chatterjee, Kourosh Saeb Parsy, Anthony R. Green, Jyoti Nangalia, Elisa Laurenti, and Peter J. Campbell. Clonal dynamics of haematopoiesis across the human lifespan. Nature, 606(7913):343–350, June 2022. ISSN 1476-4687. doi: 10.1038/s41586-022-04786-y. URL https://www.nature.com/articles/s41586-022-04786-y. Publisher: Nature Publishing Group.

Marcus T Moen and Iain G Johnston. HyperHMM: efficient inference of evolutionary and progressive dynamics on hypercubic transition graphs. Bioinformatics, 39(1):btac803, January 2023. ISSN 1367-4811. doi: 10.1093/bioinformatics/btac803. URL https://doi.org/10.1093/bioinformatics/btac803.

Claude Monnet, Kenneth De Baets, and Christian Klug. Parallel evolution controlled by adaptation and covariation in ammonoid cephalopods. BMC Evolutionary Biology, 11(1):115, December 2011. ISSN 1471-2148. doi: 10.1186/1471-2148-11-115. URL http://bmcevolbiol.biomedcentral.com/articles/10.1186/1471-2148-11-115.

Martha M. Muñoz, Luke O. Frishkoff, Jenna Pruett, and D. Luke Mahler. Evolution of a Model System: New Insights from the Study of Anolis Lizards. Annual Review of Ecology, Evolution and Systematics, 54(Volume 54, 2023):475–503, November 2023. ISSN 1543-592X, 1545-2069. doi: 10.1146/annurev-ecolsys-110421-103306. URL https://www.annualreviews.org/content/journals/10.1146/annurev-ecolsys-110421-103306. Publisher: Annual Reviews.

Mark Pagel. The Maximum Likelihood Approach to Reconstructing Ancestral Character States of Discrete Characters on Phylogenies. Systematic Biology, 48(3):612–622, 1999. ISSN 1063-5157. URL https://www.jstor.org/stable/2585328. Publisher: [Oxford University Press, Society of Systematic Biologists].

Steven Poe, Adrían Nieto-montes De Oca, Omar Torres-carvajal, Kevin De Queiroz, Julían A. Velasco, Brad Truett, Levi N. Gray, Mason J. Ryan, Gunther Köhler, Fernando Ayala-varela, and Ian Latella. A Phylogenetic, Biogeographic, and Taxonomic study of all Extant Species of Anolis (Squamata; Iguanidae). Systematic Biology, 66(5):663–697, September 2017. ISSN 1063-5157, 1076-836X. doi: 10.1093/sysbio/syx029. URL http://academic.oup.com/sysbio/article/66/5/663/3056323/A-Phylogenetic-Biogeographic-and-Taxonomic-study.

Mikael Pontarp. Ecological opportunity and adaptive radiations reveal eco-evolutionary perspectives on community structure in competitive communities. Scientific Reports, 11(1): 19560, October 2021. ISSN 2045-2322. doi: 10.1038/s41598-021-98842-8. URL https://www.nature.com/articles/s41598-021-98842-8. Number: 1 Publisher: Nature Publishing Group.

M. N. Puttick. Mixed evidence for early bursts of morphological evolution in extant clades. Journal of Evolutionary Biology, 31(4):502–515, 2018. ISSN 1420-9101. doi: 10.1111/jeb.13236. URL https://onlinelibrary.wiley.com/doi/abs/10.1111/jeb.13236. eprint: https://onlinelibrary.wiley.com/doi/pdf/10.1111/jeb.13236.

Liam J. Revell. phytools: an R package for phylogenetic comparative biology (and other things): *phytools: R package*. Methods in Ecology and Evolution, 3(2):217–223, April 2012. ISSN 2041210X. doi: 10.1111/j.2041-210X.2011.00169.x. URL https://onlinelibrary.wiley.com/doi/10.1111/j.2041-210X.2011.00169.x.

Liam J. Revell. phytools 2.0: an updated R ecosystem for phylogenetic comparative methods (and other things). PeerJ, 12:e16505, January 2024. ISSN 2167-8359. doi: 10.7717/peerj.16505. URL https://pmc.ncbi.nlm.nih.gov/articles/PMC10773453/.

Nicolas Rodrigue, Hervé Philippe, and Nicolas Lartillot. Uniformization for sampling realizations of Markov processes: applications to Bayesian implementations of codon substitution models. Bioinformatics, 24(1):56–62, January 2008. ISSN 1367-4803. doi: 10.1093/bioinformatics/btm532. URL https://doi.org/10.1093/bioinformatics/btm532.

Rebecca J. Rundell and Trevor D. Price. Adaptive radiation, nonadaptive radiation, ecological speciation and nonecological speciation. Trends in Ecology & Evolution, 24(7):394–399, July 2009. ISSN 0169-5347. doi: 10.1016/j.tree.2009.02.007. URL https://www.sciencedirect.com/science/article/pii/S0169534709001268.

Michael J. Sanderson and Michael J. Donoghue. Patterns of Variation in Levels of Homoplasy. Evolution, 43(8):1781–1795, 1989. ISSN 1558-5646. doi: 10.1111/j.1558-5646.1989.tb02626.x. URL https://onlinelibrary.wiley.com/doi/abs/10.1111/j.1558-5646.1989.tb02626.x. eprint: https://onlinelibrary.wiley.com/doi/pdf/10.1111/j.1558-5646.1989.tb02626.x.

Hervé Sauquet, Maria von Balthazar, Susana Magalĺon, James A. Doyle, Peter K. Endress, Emily J. Bailes, Erica Barroso de Morais, Kester Bull-Hereñu, Laetitia Carrive, Marion Chartier, Guillaume Chomicki, Mario Coiro, Raphäel Cornette, Juliana H. L. El Ottra, Cyril Epicoco, Charles S. P. Foster, Florian Jabbour, Agathe Haevermans, Thomas Haevermans, Rebeca Herńandez, Stefan A. Little, Stefan Löfstrand, Javier A. Luna, Julien Massoni, Sophie Nadot, Susanne Pamperl, Charlotte Prieu, Elisabeth Reyes, Patŕıcia dos Santos, Kristel M. Schoonderwoerd, Susanne Sontag, Anäelle Soulebeau, Yannick Staedler, Georg F. Tschan, Amy Wing-Sze Leung, and Jürg Schönenberger. The ancestral flower of angiosperms and its early diversification. Nature Communications, 8(1):16047, August 2017. ISSN 2041-1723. doi: 10.1038/ncomms16047. URL https://www.nature.com/articles/ncomms16047. Publisher: Nature Publishing Group.

Orlando Schwery, Will Freyman, and Emma E. Goldberg. adequaSSE: Model Adequacy Testing for Trait-Dependent Diversification Models. preprint, Evolutionary Biology, March 2023. URL http://biorxiv.org/lookup/doi/10.1101/2023.03.06.531416.

Alex Skeels. Lineages through space and time plots: Visualising spatial and temporal changes in diversity. Frontiers of Biogeography, 11(2), 2019. doi: 10.21425/F5FBG42954. URL https://escholarship.org/uc/item/1521z9pb.

Tanja Stadler. Lineages-through-time plots of neutral models for speciation. Mathematical Biosciences, 216(2):163–171, December 2008. ISSN 00255564. doi: 10.1016/j.mbs.2008.09.006. URL https://linkinghub.elsevier.com/retrieve/pii/S0025556408001557.

James T. Stroud and Jonathan B. Losos. Ecological Opportunity and Adaptive Radiation. Annual Review of Ecology, Evolution, and Systematics, 47(1):507–532, November 2016. ISSN 1543-592X, 1545-2069. doi: 10.1146/annurev-ecolsys-121415-032254. URL https://www.annualreviews.org/doi/10.1146/annurev-ecolsys-121415-032254.

James T Stroud and Jonathan B Losos. Bridging the Process-Pattern Divide to Understand the Origins and Early Stages of Adaptive Radiation: A Review of Approaches With Insights From Studies of Anolis Lizards. Journal of Heredity, 111(1):33–42, February 2020. ISSN 0022-1503, 1465-7333. doi: 10.1093/jhered/esz055. URL https://academic.oup.com/jhered/article/111/1/33/5644556.

Sergei Tarasov, Istváan Mikáo, Matthew Jon Yoder, and Josef C Uyeda. PARAMO: A Pipeline for Reconstructing Ancestral Anatomies Using Ontologies and Stochastic Mapping. Insect Systematics and Diversity, 3(6):1, November 2019. ISSN 2399-3421. doi: 10.1093/isd/ixz009. URL https://doi.org/10.1093/isd/ixz009.

Evan Twomey, Paulo Melo-Sampaio, Lisa M Schulte, Franky Bossuyt, Jason L Brown, and Santiago Castroviejo-Fisher. Multiple Routes to Color Convergence in a Radiation of Neotropical Poison Frogs. Systematic Biology, 72(6):1247–1261, November 2023. ISSN 1063-5157. doi: 10.1093/sysbio/syad051. URL https://doi.org/10.1093/sysbio/syad051.

Julían A. Velasco, Fabricio Villalobos, Jośe A. F. Diniz-Filho, Steven Poe, and Oscar Flores-Villela. Macroecology and macroevolution of body size in Anolis lizards. Ecography, 43 (6):812–822, 2020. ISSN 1600-0587. doi: 10.1111/ecog.04583. URL https://onlinelibrary.wiley.com/doi/abs/10.1111/ecog.04583. eprint: https://nsojournals.onlinelibrary.wiley.com/doi/pdf/10.1111/ecog.04583.

Roberto Vendramin, Kevin Litchfield, and Charles Swanton. Cancer evolution: Darwin and beyond. The EMBO Journal, 40(18):e108389, September 2021. ISSN 0261-4189. doi: 10.15252/embj.2021108389. URL https://www.embopress.org/doi/full/10.15252/embj.2021108389. Publisher: John Wiley & Sons, Ltd.

Ben P Williams, Iain G Johnston, Sarah Covshoff, and Julian M Hibberd. Phenotypic landscape inference reveals multiple evolutionary paths to C4 photosynthesis. eLife, 2:e00961, September 2013. ISSN 2050-084X. doi: 10.7554/eLife.00961. URL https://doi.org/10.7554/eLife.00961. Publisher: eLife Sciences Publications, Ltd.

Michael L. Yuan, Marvalee H. Wake, and Ian J. Wang. Phenotypic integration between claw and toepad traits promotes microhabitat specialization in the *Anolis* adaptive radiation. Evolution, 73(2):231–244, February 2019. ISSN 0014-3820, 1558-5646. doi: 10.1111/evo.13673. URL https://academic.oup.com/evolut/article/73/2/231/6882169.

Walter Zimmermann. Research on Phylogeny of Species and of Single Characters. The American Naturalist, 68(717):381–384, 1934. ISSN 0003-0147. URL https://www.jstor.org/stable/2456939. Publisher: [The University of Chicago Press, The American Society of Naturalists].

Walter Zimmermann. Main results of the “Telome Theory”. Journal of Palaeosciences, 1: 456–470, December 1952. ISSN 2583-4266. doi: 10.54991/jop.1952.423. URL https://www.jpsonline.co.in/index.php/jop/article/view/423.

