## Supplement for "Summarizing Evolutionary Trajectories from Phylogenetic Character Maps of Discrete Traits"

Version dated: June 14, 2026

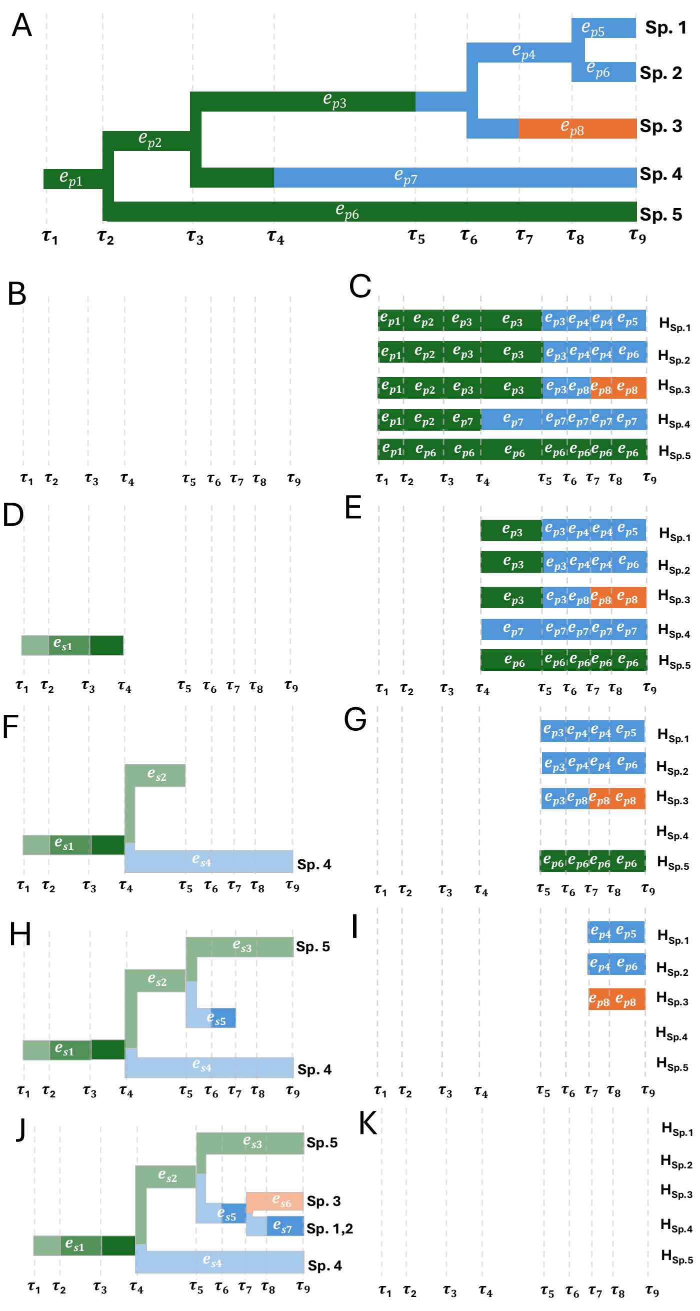


Figure 1: Scenario tree construction cartoon.

Figure 1: Scenario tree construction cartoon. The scenario tree compression algorithm requires (A) a phylogenetic tree with discrete character states (orange, blue, and green) mapped along it. Edges are labelled $e_{p,i}$. Time is discretized into a shared vector of event times $\tau = (\tau_1, \ldots, \tau_9)$, spanning from the root ($\tau_1$) to the present ($\tau_9$). The algorithm is initialized with (B) an empty scenario tree object with the same global set of time slices $\tau$, defined by all event times in the tree. As the scenario tree is constructed, edges labelled $e_{s,i}$ will be added to the tree; and (C) a matrix representation of lineage histories through time, where each row corresponds to a species and each column to a time slice, with entries tracking the active phylogenetic edge and its associated state. (D)-(K) illustrate scenario construction: starting at the root, all lineages are grouped into a single scenario edge, and as time advances, lineages that remain in the same state and follow the same phylogenetic segment remain grouped; when a state transition occurs, the scenario splits, with one branch continuing the original state and a second representing transitioning lineages; at speciation events, descendant lineages inherit the current grouping unless they differ in state.

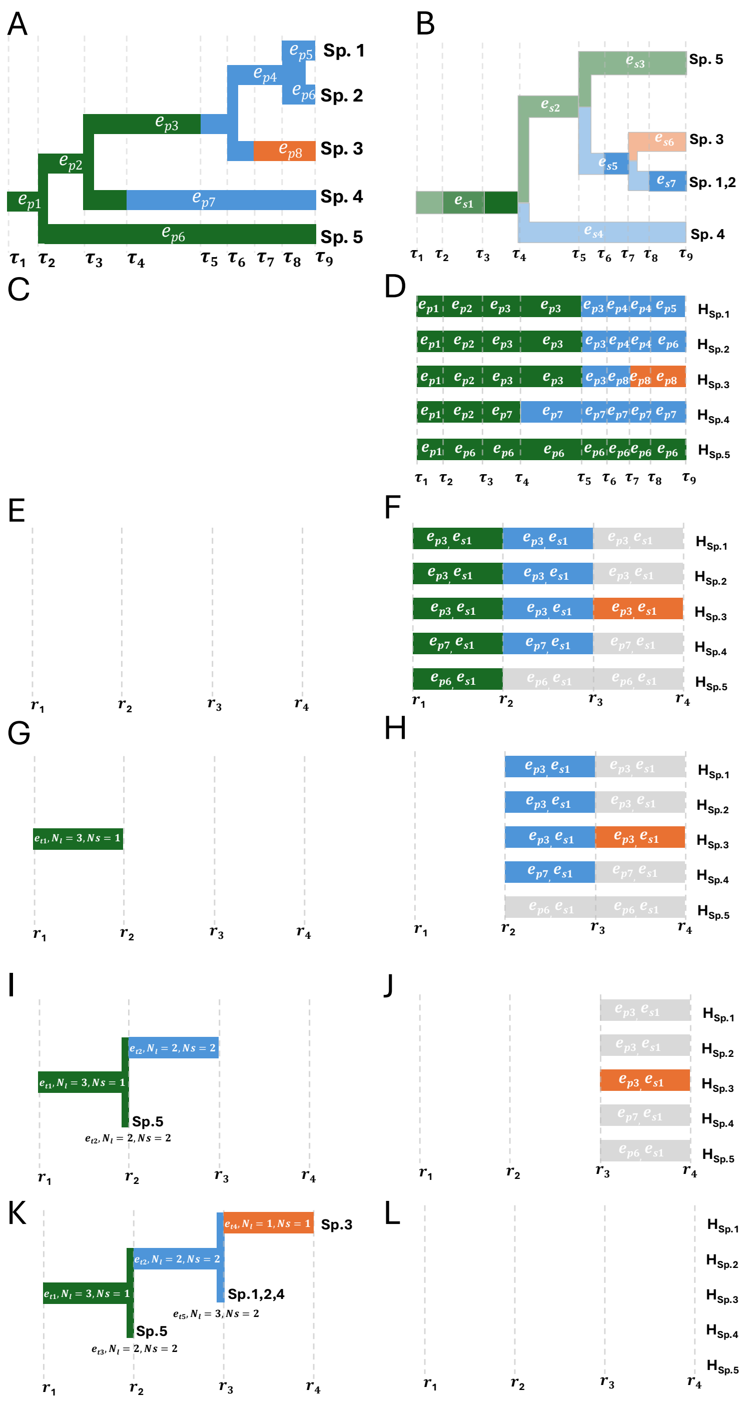


Figure 2: Scenario tree construction cartoon.

Figure 2: Transition tree construction cartoon. The transition tree compression algorithm requires (A) a phylogenetic tree with discrete character states (orange, blue, and green) mapped along it. Edges are labelled $e_{p,i}$. (B) a scenario tree with discrete character states (orange, blue, and green) mapped along it. Edges are labelled $e_{s,i}$. Time is discretized into a shared vector of event times $\tau = (\tau_1, \ldots, \tau_9)$, spanning from the root ($\tau_1$) to the present ($\tau_9$). The algorithm is initialized with (C) an empty scenario tree object with branch lengths in units of unique state transitions $r$. As the transition tree is constructed, edges labelled $e_{t,i}$ will be added to the tree each including the number of lineages and scenarios associated with it; and (D) a matrix representation of lineage transition sequences, where each row corresponds to a species and each column to a transition sequence step, with entries tracking the active phylogenetic edge and its associated state. This is a major difference from scenario tree construction, where the matrix is sorted into time steps not transition sequence steps. (D)-(K) illustrate transition tree construction: starting at the root and first transition sequence step shared by all species (in this case green), all lineages are grouped into a single transition tree edge. We record the number of lineages and scenarios that follow this transition sequence step. The root transition tree edge will always have one scenario associated with and 1 or more lineages. We then check if any lineages have a second step in their transition sequence; if so, the transition tree undergoes a splitting event. Lineages remaining within the ancestral state until their termination (present or extinction) remain in a new daughter lineage, the same color as the parent state, but a length of 0.0 (since it is not a transition sequence step). Lineages that follow an a new transition sequence step follow new transition sequence branches of length 1.0. This splitting process is repeated until the longest transition sequence is reached.

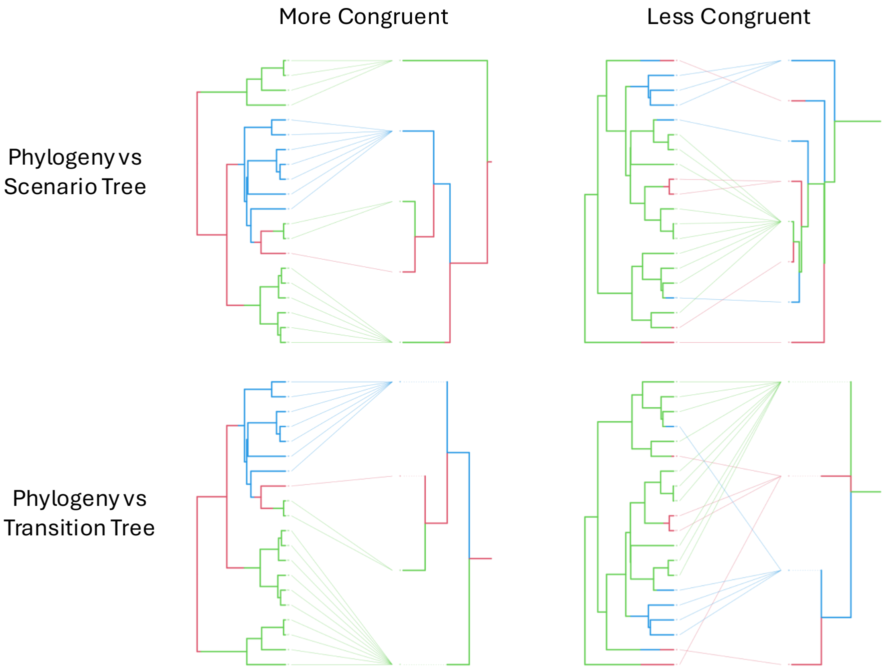


Figure 3: Example tanglegrams between phylogenetic character histories and corresponding scenario or transition trees. Examples provided show instances where scenario/transition trees closely match the phylogenetic character history, in which scenario or transition sequence divergence events (referred to as lineage-differentiating and path-unique state transitions, respectively) occur successively within a clade and are closely coupled to lineage divergence events. Such tanglegrams include a few "tangles" or crossing lines linking tips of phylogenies and scenario/transition trees. Other examples provided show instances where scenario/transition trees are not congruent with the phylogenetic character history, where scenario or transition sequence divergence events accumulate less frequently within a clade, and many clades are differentiated from others by only a single transition from an ancestral state.

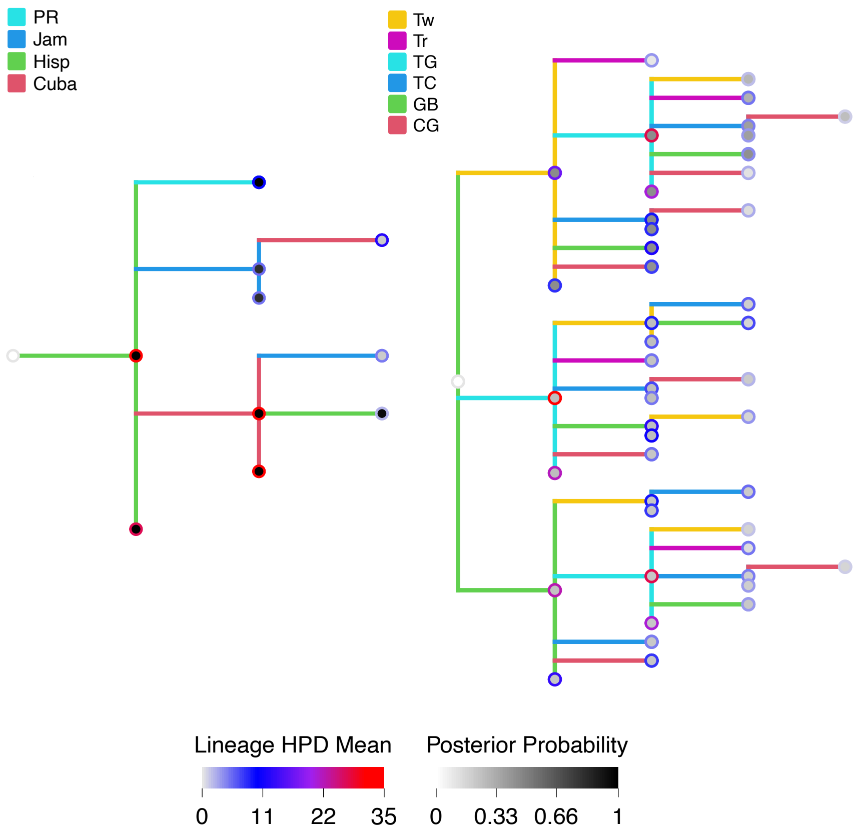


Figure 4: *Anolis* transition trees for ecomorph and regions. Transition trees summarize transition sequences across the distribution of 100 stochastic character maps. Node markers show the mean number of lineages (inner) that passed through at given transition sequence step, and mean number of lineages within a given transition sequence. Tip markers indicate the number of lineages that terminate at a given transition sequence step.

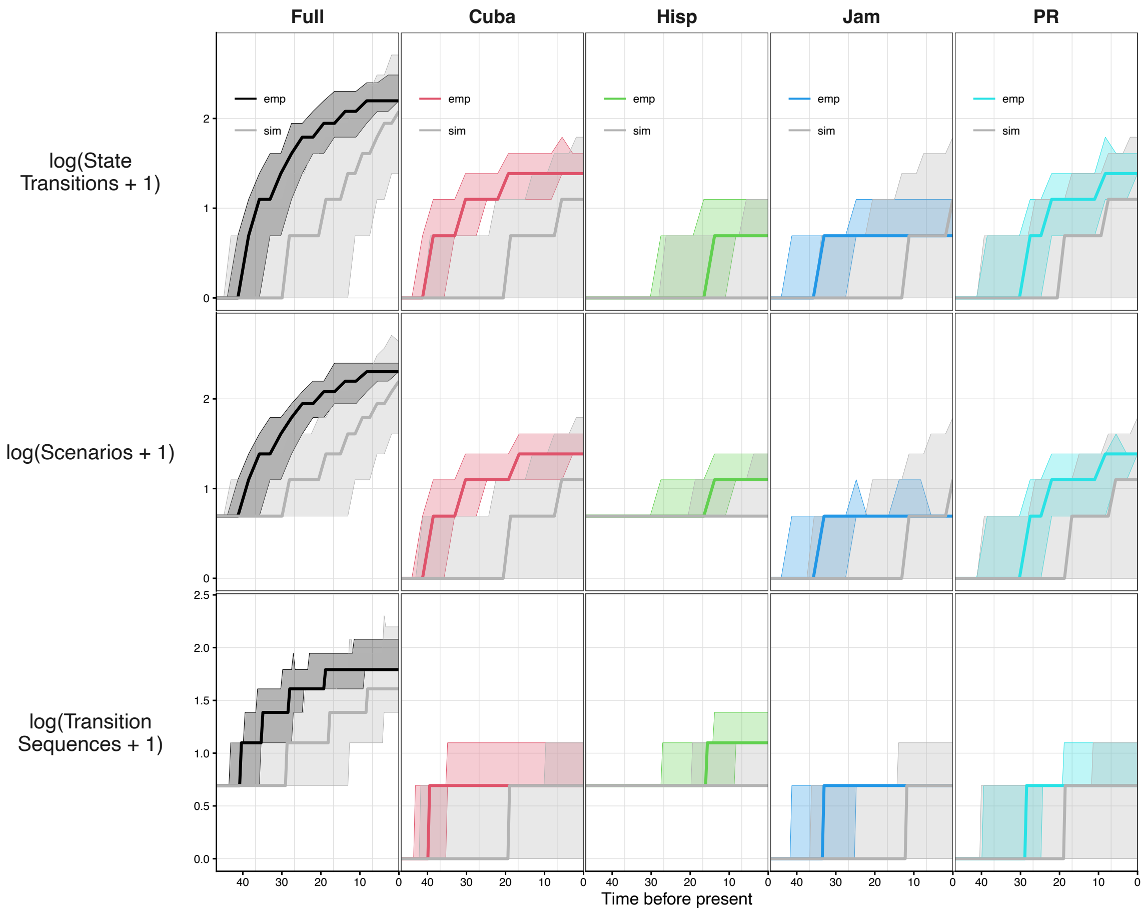


Figure 5: *Anolis* through-time distribution plots from the region statspace stochastic character maps for states transition, scenarios, and transition sequences. Transparent colors show 95% intervals while solid lines show distribution means. Each plot includes two sets of through-time distributions: an empirical distribution of stochastic character maps given the best-fit model and observed tip states (black and colored lines), and a simulated distribution (grey). Simulated distributions were generated by sampling 100 replicates of tip data from the best-fit model and the *Anolis* phylogeny, then refitting the simulated data to the best-fit model, yielding a distribution of 100 simmaps for each simulated dataset. Columns display either the "full" number or the number for the indicated region or ecomorph. For region-specific columns, state transitions count the number of transitions into states that include the specified region.

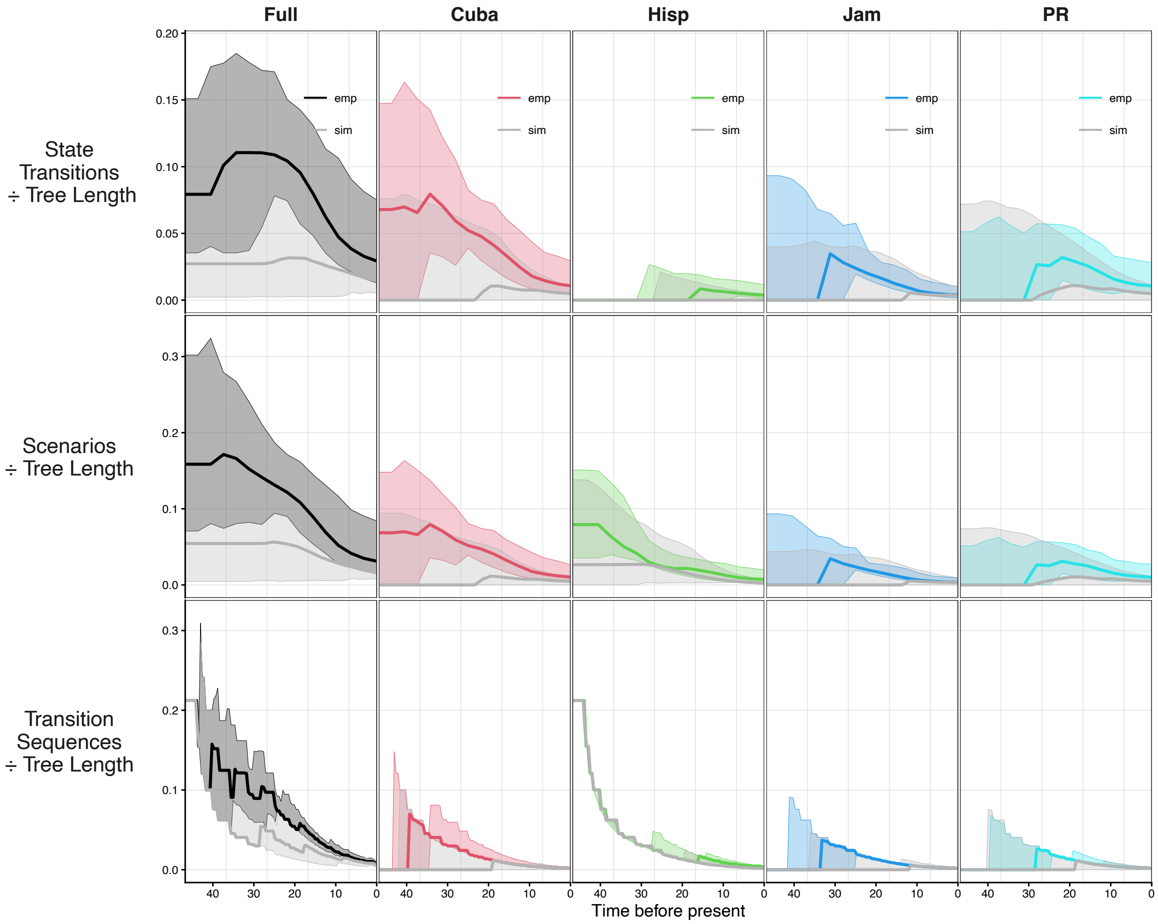


Figure 6: *Anolis* tree length normalized through-time distribution plots from the region statspace stochastic character maps for states transition, scenarios, and transition sequences. Transparent colors show 95% intervals while solid lines show distribution means. Each plot includes two sets of through-time distributions: an empirical distribution of stochastic character maps given the best-fit model and observed tip states (black and colored lines), and a simulated distribution (grey). Simulated distributions were generated by sampling 100 replicates of tip data from the best-fit model and the *Anolis* phylogeny, then refitting the simulated data to the best-fit model, yielding a distribution of 100 simmaps for each simulated dataset. Columns display either the "full" number or the number for the indicated region or ecomorph. For region-specific columns, state transitions count the number of transitions into states that include the specified region.

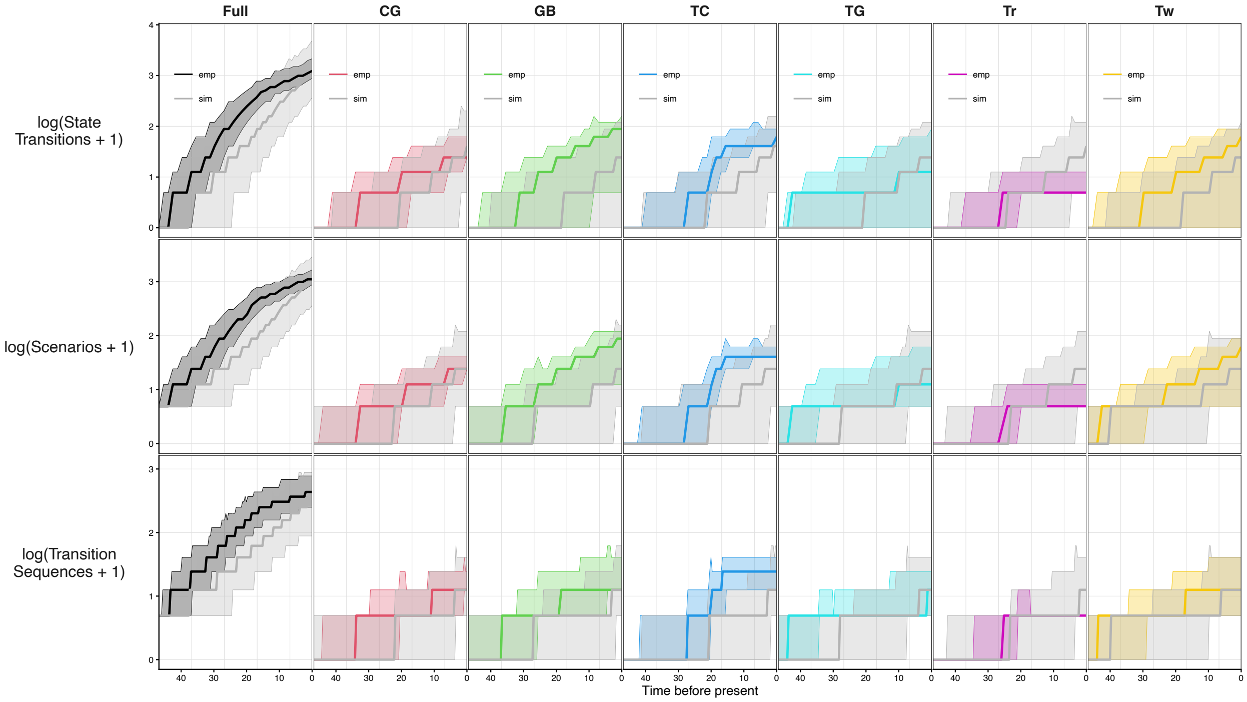


Figure 7: *Anolis* through-time distribution plots from the region statspace stochastic character maps for states transition, scenarios, and transition sequences. Transparent colors show 95% intervals while solid lines show distribution means. Each plot includes two sets of through-time distributions: an empirical distribution of stochastic character maps given the best-fit model and observed tip states (black and colored lines), and a simulated distribution (grey). Simulated distributions were generated by sampling 100 replicates of tip data from the best-fit model and the *Anolis* phylogeny, then refitting the simulated data to the best-fit model, yielding a distribution of 100 simmaps for each simulated dataset. Columns display either the "full" number or the number for the indicated region or ecomorph. For region-specific columns, state transitions count the number of transitions into states that include the specified region.

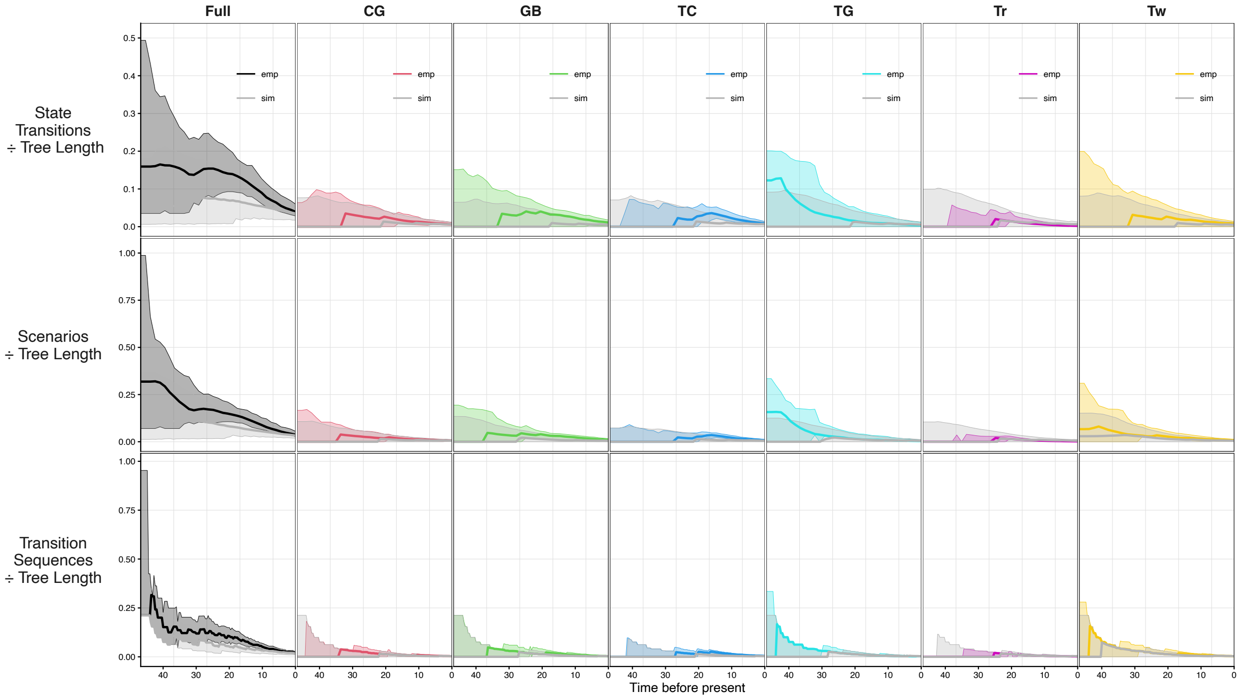


Figure 8: *Anolis* tree length normalized through-time distribution plots from the ecomorph statspace stochastic character maps for states transition, scenarios, and transition sequences. Transparent colors show 95% intervals while solid lines show distribution means. Each plot includes two sets of through-time distributions: an empirical distribution of stochastic character maps given the best-fit model and observed tip states (black and colored lines), and a simulated distribution (grey). Simulated distributions were generated by sampling 100 replicates of tip data from the best-fit model and the *Anolis* phylogeny, then refitting the simulated data to the best-fit model, yielding a distribution of 100 simmaps for each simulated dataset. Columns display either the "full" number or the number for the indicated region or ecomorph. For ecomorph-specific columns, state transitions count the number of transitions into states that include the specified region.

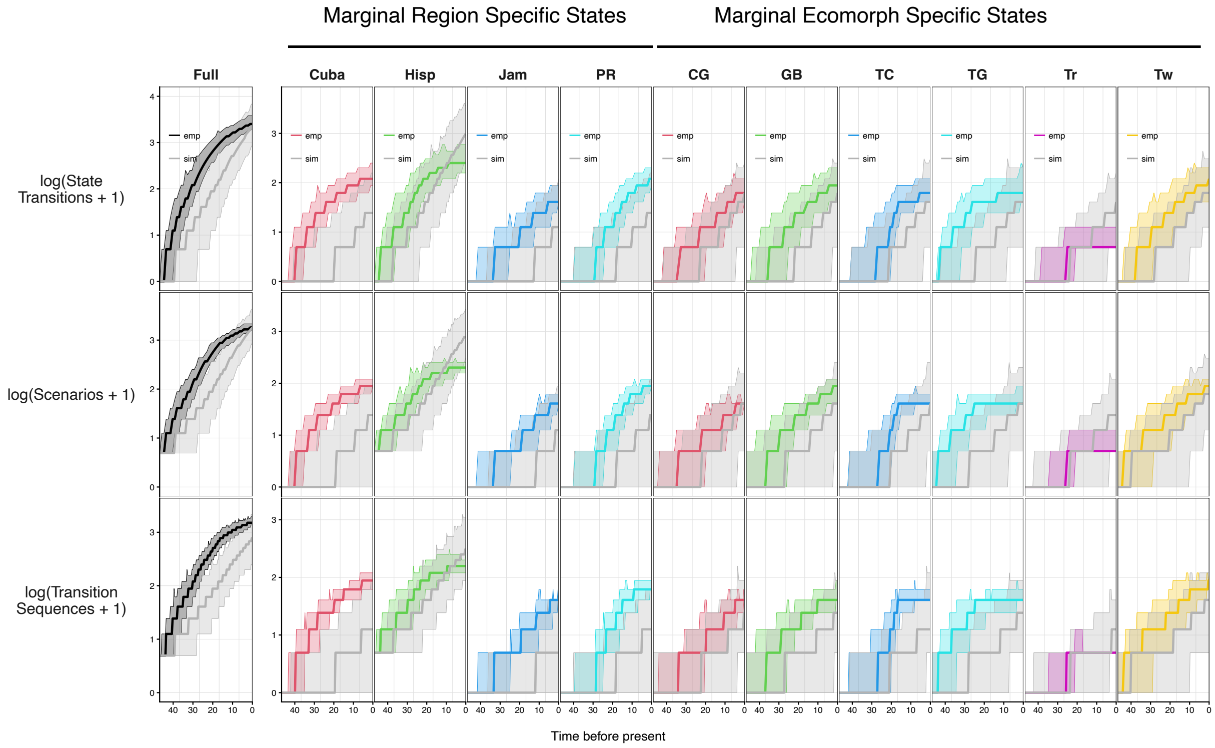


Figure 9: *Anolis* through-time distribution plots from the combined region and ecomorph statspace stochastic character maps for lineages, states transition, scenarios, and transition sequences. Transparent colors show 95% intervals while solid lines show distribution means. Each plot includes two sets of through-time distributions: an empirical distribution of stochastic character maps given the best-fit model and observed tip states (black and colored lines), and a simulated distribution (grey). Simulated distributions were generated by sampling 100 replicates of tip data from the best-fit model and the *Anolis* phylogeny, then refitting the simulated data to the best-fit model, yielding a distribution of 100 simmaps for each simulated dataset. Columns display either the ”full” number or the number for the indicated region or ecomorph. For region- and ecomorph-specific columns, state transitions count the number of transitions into states that include the specified region or ecomorph.

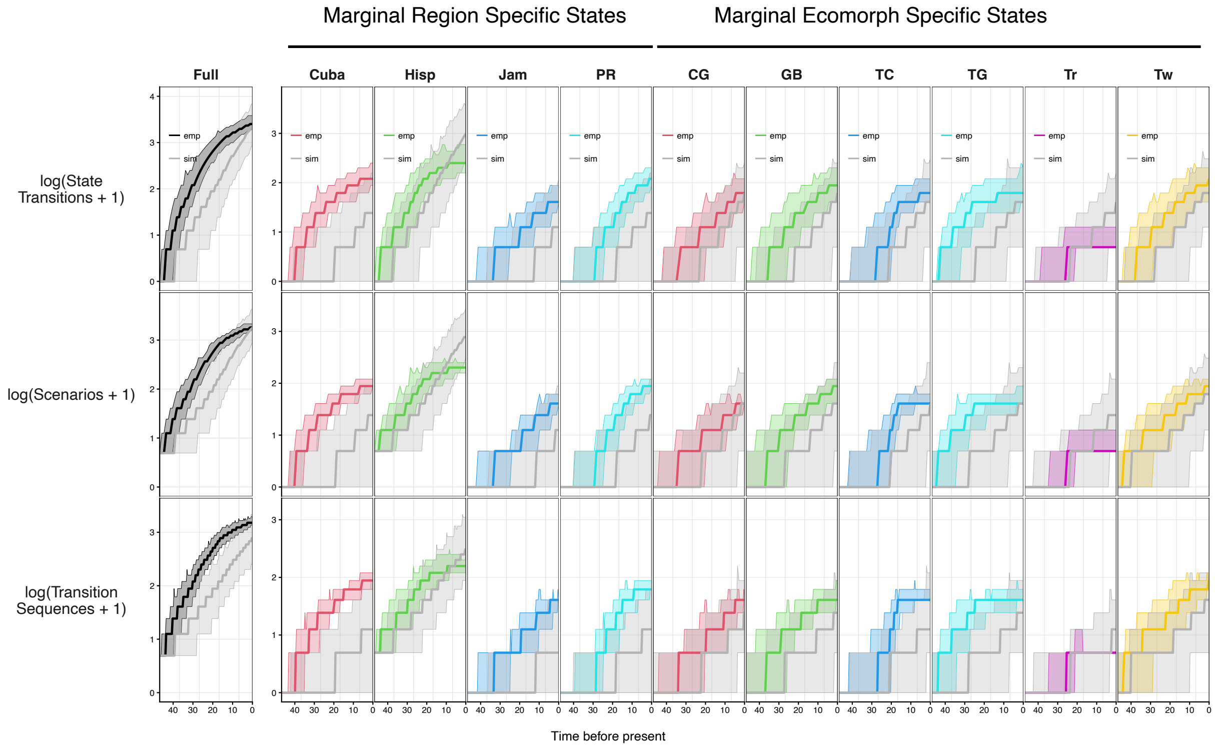


Figure 10: *Anolis* tree length normalized hrough-time distribution plots from the combined region and ecomorph statspace stochastic character maps for lineages, states transition, scenarios, and transition sequences. Transparent colors show 95% intervals while solid lines show distribution means. Each plot includes two sets of through-time distributions: an empirical distribution of stochastic character maps given the best-fit model and observed tip states (black and colored lines), and a simulated distribution (grey). Simulated distributions were generated by sampling 100 replicates of tip data from the best-fit model and the *Anolis* phylogeny, then refitting the simulated data to the best-fit model, yielding a distribution of 100 simmaps for each simulated dataset. Columns display either the "full" number or the number for the indicated region or ecomorph. For region- and ecomorph-specific columns, state transitions count the number of transitions into states that include the specified region or ecomorph.
